# Lipid remodelling enables adaptation to chronic hyperosmotic stress

**DOI:** 10.64898/2026.03.26.714418

**Authors:** Thomas D. Williams, Christian Klose, Robert Ernst, Pedro Carvalho

## Abstract

Lipid droplets (LDs) accumulate in response to diverse cellular stresses. However, their regulation and physiological roles remain poorly understood in most contexts. Here, we show that, in budding yeast, chronic hyperosmotic stress induces sustained LD accumulation. Unlike the transient LD response observed during acute osmotic shock, chronic stress triggers prolonged, Dga1-dependent triacylglycerol synthesis. In the absence of triacylglycerol synthesis cellular fitness is severely affected. Lipidomic profiling reveals extensive membrane remodelling during chronic hyperosmotic stress, most notably a shift from phosphatidylethanolamine to phosphatidylcholine. In LD-deficient cells, the stress–induced PC increase is blunted and manipulation of PC synthesis modulates the fitness of triacylglycerol-deficient cells under hyperosmotic stress. Thus, LD accumulation and phospholipid remodelling underlie an adaptive response to chronic hyperosmotic stress.

**Summary:** This work demonstrates that membrane remodelling occurs in cells experiencing chronic hyperosmotic stress. Both triacylglycerol and phosphatidylcholine levels are increased. Cell fitness depends upon increased triacylglycerol synthesis and is further modulated by manipulating phosphatidylcholine levels.

## Introduction

Lipid droplets (LDs) are unique lipid storage organelles present in most eukaryotic cells (Mathiowetz and Olzmann, 2024; Farese and Walther, 2025; Klemm and Carvalho, 2024). They are composed of a phospholipid monolayer surrounding a hydrophobic core of neutral lipids, primarily triacylglycerols (TAGs) and sterol esters (STEs). TAGs are synthesized through the acylation of diacylglycerol (DAG) with a fatty acid, either derived from an activated acyl–CoA or transferred from a donor phospholipid (Athenstaedt, 2010). STEs are generated by esterifying sterols with acylated fatty acids (Athenstaedt, 2010). LD biogenesis occurs by an evolutionarily conserved mechanism where a Seipin oligomer concentrates the neutral lipids within the ER bilayer, thereby facilitating their condensation into lens structures that subsequently bud towards the cytoplasm to originate a mature LD (Klug et al., 2024; Thiam and Ikonen, 2021).

Proper LD assembly and regulation are essential for cellular and organismal health through membrane property optimisation and energy homeostasis (Zadoorian et al., 2023). Defects in LD formation result in lipodystrophies, best exemplified by the seipinopathies (Cartwright and Goodman, 2012), while excessive LD formation is the defining feature of fatty liver disease (Heeren and Scheja, 2021) and is further associated with poor cancer prognosis (Quan et al., 2025). When cells are challenged by environmental changes, or high stress environments such as those found in tumours, LD accumulation is widely observed (Pressly et al., 2022; Jin et al., 2023; Safi et al., 2024). Lipids stored in LDs contribute to ATP synthesis and cell adaptation to nutrient limitation (Seo et al., 2017; Rambold et al., 2015; Hariri et al., 2018; Rogers et al., 2021). When membrane expansion is required, such as upon growth resumption from quiescence, LDs can be rapidly turned over to supply lipids for membrane biogenesis (Markgraf et al., 2014). Conversely, recent evidence shows that LDs can buffer excess lipids and membranes (Phan et al., 2025; Wang et al., 2018) or sequester potentially harmful membrane species, such as those which are susceptible to oxidation (Lange et al., 2025) or which can otherwise harm organelle integrity (Nguyen and Olzmann, 2017).

Fluctuations in temperature alter membrane properties (Ernst et al., 2016), and the inability to make LDs renders cells sensitive to changes in temperature (Wang et al., 2018). While LD accumulation has been associated with various stress responses (Seo et al., 2017; Lange et al., 2025; Phan et al., 2025; Hariri et al., 2019; Lee et al., 2013; Nagaraj et al., 2023; Fei et al., 2011), these findings remain context-specific, and a comprehensive understanding of LD dynamics and regulation during stress is still missing.

Here, building on a high-throughput approach to quantify LD abundance in budding yeast, we identified a previously unrecognized role for LDs during chronic hyperosmotic stress. We show LD accumulation occurs in parallel to the activity of the canonical hyperosmotic adaptation machinery and enhances a crucial increase in phosphatidylcholine levels. Manipulating phosphatidylcholine levels affects the requirement for LD accumulation upon chronic hyperosmotic stress. Our findings demonstrate lipid remodelling is an important part of the adaptive response to chronic hyperosmotic stress.

## Results

Pln1 localizes to the surface of LDs and is rapidly degraded when not bound to LDs (Gao et al., 2017). Fluorescently tagged Pln1 is therefore commonly used as an LD marker (Figure S1A) (Gao et al., 2017; Giménez-Andrés et al., 2021; Wang et al., 2024) and its levels are a good surrogate for LD abundance (Rao et al., 2025; Gao et al., 2017). To investigate LD accumulation dynamics in a high-throughput manner, we measure the levels of endogenously tagged Pln1-mCherry using flow cytometry. During exponential growth, Pln1-mCherry levels sharply increase as cells approach stationary phase, mirroring LD abundance as detected by fluorescence microscopy of BODIPY stained cells (Figure S1A,B).

To further validate the flow cytometry-based results, we used pharmacological inhibition of TORC1 with rapamycin, a treatment known to induce LD accumulation (Madeira et al., 2015; Teixeira et al., 2021; Liu et al., 2019; Hosios et al., 2022; Dubots et al., 2014). As expected, rapamycin treatment of WT cells resulted in a rapid and sustained increase of Pln1-mCherry levels (Figure S1C). Therefore, flow cytometry analysis of Pln1-mCherry levels provides a sensitive and accurate readout of LD accumulation.

### Chronic hyperosmotic stress induces lipid droplet accumulation

We next monitored the levels of Pln1-mCherry over time in exponentially growing cells subjected to a variety of stresses: oxidative, temperature, nutrient, and hyperosmotic. Upon moderate temperature shifts (from 30°C to 25°C or 37°C) or acute restriction of glucose or amino acids (2% to 0.1% glucose or a 10-fold reduction in amino acid levels) cells showed minimal changes in Pln1-mCherry levels over a 6-hour period (Figure S1D,E). Oxidative stress induced by H2O2 resulted in a mild and gradual Pln1-mCherry accumulation (Figure S1F). In contrast, hyperosmotic stress induced by either KCl or Sorbitol resulted in a rapid increase in Pln1-mCherry accumulation that peaked at ∼3-4 hours after stress induction and remained high for at least 24 hours (Figure 1A,B and S1G,H). Moreover, the increase in Pln1-mCherry levels triggered by KCl-induced hyperosmotic stress was dose and time dependent (Figure S2A) and occurred in both rich and defined media (Figure S2B). A reduction of Pln1-mCherry to baseline levels was observed within 3 hours upon removal of the hyperosmotic stress (Figure 1C). Finally, we confirmed that the increase in Pln1-mCherry levels upon hyperosmotic stress was due to LD accumulation by imaging of BODIPY-labelled cells (Figure 1D). This analysis showed that KCl treatment increased both the number and size of LDs in cells (Figure 1E,F). Together, these results demonstrate that chronic hyperosmotic stress triggers long-lasting LD accumulation.

**Figure 1:**
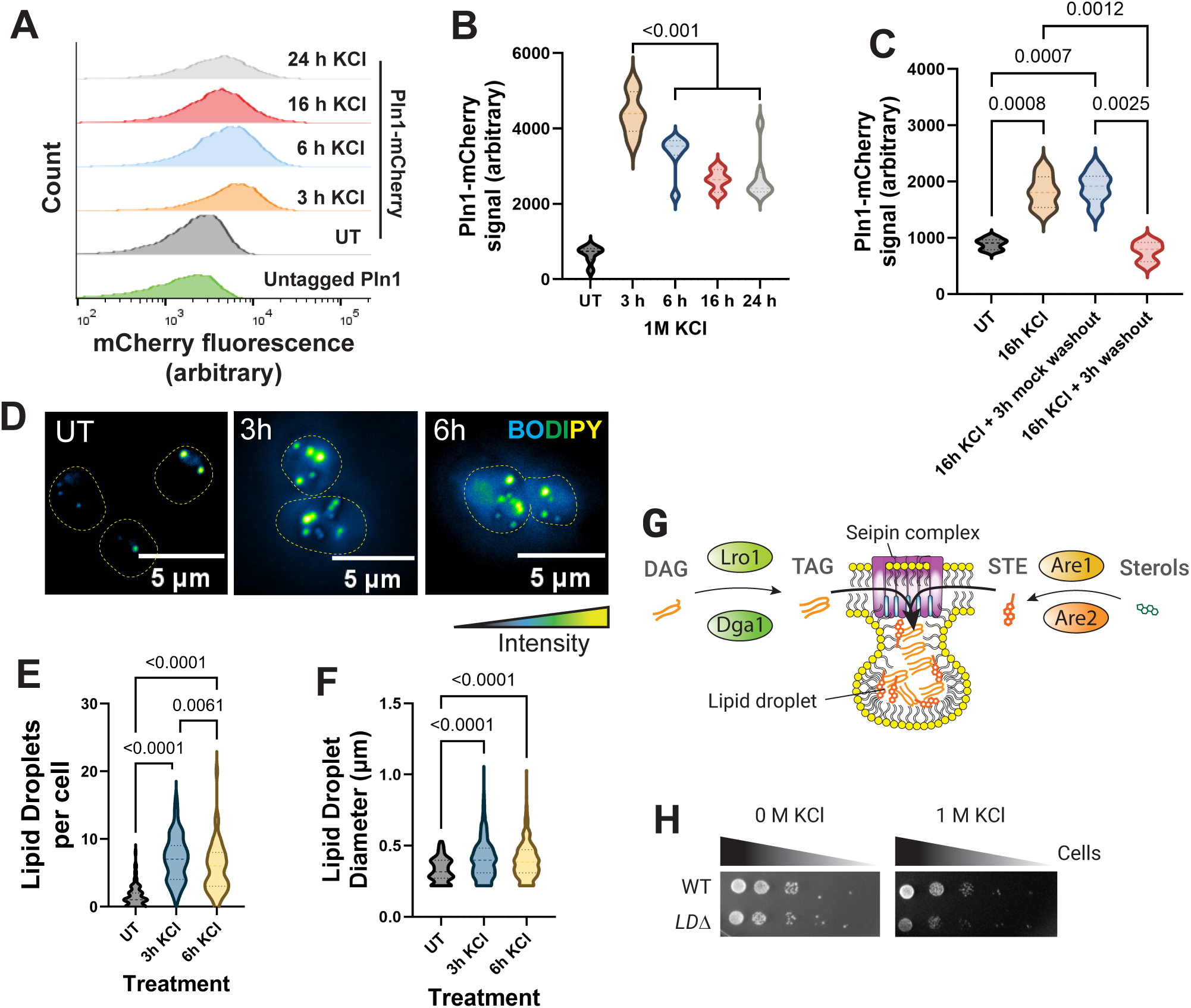
Lipid droplet accumulation is a chronic response to hyperosmotic stress. **A)** Representative histograms obtained from Pln1-mCherry tagged cells grown in SCD + 1M KCl for the indicated timepoints. The untagged Pln1 shows the background fluorescence, while UT is untreated. **B)** Median Pln1-mCherry fluorescence measured by flow cytometry from populations of single cells subjected to 1M KCl treatment in SCD media for the indicated time. UT = untreated. **C)** Median Pln1-mCherry fluorescence measured by flow cytometry from populations of single cells subjected to 16 hours of 1M KCl treatment in SCD media which was then washed out and replaced with SCD + 1M KCl (mock washout) or SCD without 1M KCl for 3 hours before measurement. UT = untreated. **D)** Lipid droplets stained with BODIPY and imaged by live cell microscopy after varying lengths of 1M KCl treatment in SCD. Z-projection images representative of multiple replicates. UT = untreated. **E)** Analysis of lipid droplets per cell from samples used in (C). UT = untreated. **F)** Analysis of lipid droplet diameters from samples used in (C). UT = untreated. **G)** Schematic of lipid droplet formation. Dga1 or Lro1 converts diacylglycerol (DAG) into triacylglycerol (TAG), while Are1 or Are2 convert sterols into sterol esters (STE). TAGs and STEs are concentrated and packaged into lipid droplets by the Seipin complex. **H)** Growth assay of wild-type (WT) and a*re1Δare2Δdga1Δlro1Δ* (*LDΔ*) cells after 2 days on SCD agar and SCD + 1M KCl agar.

### Lipid droplet accumulation is important for cell fitness under hyperosmotic stress

Hyperosmotic stress causes rapid water egress and a drop in membrane tension (Roffay et al. 2021; Phan et al. 2025). However, these effects, which impact LDs (Phan et al., 2025), are typically resolved within 30-60 minutes of stress onset (Klipp et al., 2005; Shen et al., 2023; Petelenz-Kurdziel et al., 2013), long before the phenotypes we observe. We therefore hypothesized that LD accumulation may be important for long-term fitness under hyperosmotic stress. LD biogenesis depends on neutral lipid synthesis and is spatially controlled by the seipin complex (Figure 1G). Seipin mutants produce aberrant LDs that are defective in recruiting surface proteins including Pln1-mCherry (Grippa et al., 2015). Consistent with this observation, seipin mutants showed reduced Pln1-mCherry accumulation under hyperosmotic stress (Figure S2C), and most cells had only one or two large LDs (Figure S2D,E). Since the existing LDs expanded under stress, neutral lipids were still produced (Figure S2F). Of note, the seipin dependent changes in LD size did not compromise cell fitness under hyperosmotic conditions (Figure S2G). However, when we tested the importance of stress induced neutral lipid synthesis with a strain that lacks all four relevant enzymes (Dga1 and Lro1, which produce TAGs, and Are1 and Are2, which synthesise STEs) (Oelkers et al. 2002; Sandager et al. 2002), the cells displayed a severe growth defect under hyperosmostic conditions. (Fig 1H). We therefore concluded that LD accumulation is required for adaptation to hyperosmotic stress.

### Hyperosmotic lipid droplet accumulation is driven by Dga1-mediated triacylgycerol synthesis

We next assessed the individual contribution of TAG and STE towards cell fitness under hyperosmotic stress. Simultaneous deletion of Are1 and Are2, which prevent STE synthesis, did not impact cell growth either in the absence or presence of hyperosmotic stress (Figure 2A). In contrast, preventing TAG synthesis with *dga1Δlro1Δ* cells resulted in a growth defect specifically under hyperosmotic stress (Figure 2A). Deletion of either *dga1* or *lro1* alone resulted in growth comparable to WT cells (Figure 2A). Therefore, TAG synthesis is required to adaptation to chronic hyperosmotic stress with Dga1 and Lro1 playing largely redundant roles.

**Figure 2:**
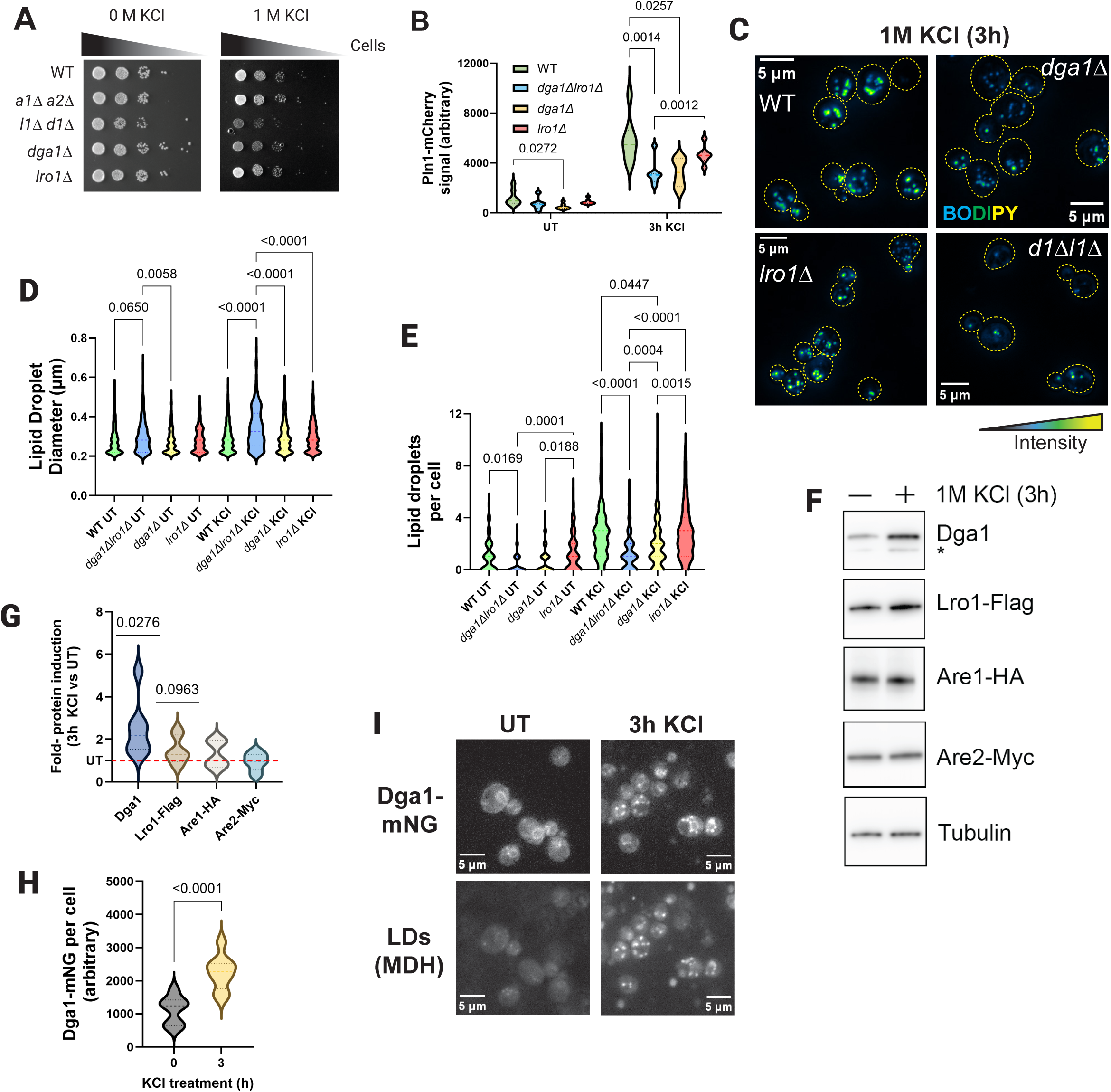
Dga1 produced triacylglycerols drive chronic hyperosmotic lipid droplet accumulation. **A)** Growth assay of wild-type (WT), a*re1Δare2Δ* (*a1Δa2Δ*), *dga1Δlro1Δ* (*d1Δl1Δ*), *dga1Δ*, and *lro1Δ* cells after 2 days on SCD agar and SCD + 1M KCl agar. **B)** Median Pln1-mCherry fluorescence measured by flow cytometry from populations of single wild-type (WT), *dga1Δlro1Δ, dga1Δ*, and *lro1Δ* cells subjected to 1M KCl treatment in SCD media for 3 hours. UT = untreated. **C)** Lipid droplets stained with BODIPY and imaged by live cell microscopy after 3 hours 1M KCl treatment in SCD for Pln1-mCherry wild-type (WT), *dga1Δlro1Δ, dga1Δ*, and *lro1Δ* cells. Z-projection images representative of multiple replicates. **D)** Analysis of lipid droplets per cell from samples used in (C). UT = untreated. **E)** Analysis of lipid droplet diameters from samples used in (C). UT = untreated. **F)** Western blot analysis of Dga1, Lro1-Flag, Are1-HA, and Are2-Myc levels before and after 3 hours of 1M KCl treatment in SCD. Representative of multiple replicates. **G)** Analysis of normalised protein fold-change from F. Statistics are compared to 1 (no fold-change). UT = untreated. **H)** Median Dga1-mNeonGreen fluorescence measured by flow cytometry from populations of cells subjected to 1M KCl treatment in SCD media for the indicated time. **I)** Visualisation of Dga1-mNeonGreen (Dga1-mNG) and lipid droplets (LD, stained using MDH) in live cells with and without 3 hours of treatment with 1M KCl in SCD. UT = untreated.

Since the growth assays reveal the importance of TAG synthesis for long-term adaptation, we monitored Pln1-mCherry to investigate LD accumulation upon hyperosmotic stress. Consistent with the growth assays, *are1Δare2Δ* cells displayed no changes in LD level compared to the WT using Pln1-mCherry or BODIPY as markers. However, the LD response was strongly blunted in *dga1Δlro1Δ* cells (Figures 2B-E, S2J-M). In contrast to acute hyperosmotic stress (Phan et al., 2025), Lro1 is dispensable for LD accumulation induced by chronic hyperosmotic stress (Figures 2B-E). On the other hand, *dga1Δ* cells showed much less Pln1-mCherry, comparable to that observed in *dga1Δlro1Δ* cells (Figure 2B). The defect in LD accumulation in *dga1Δ* cells was further confirmed by imaging LDs after BODIPY-labelling. LDs (Figure 2C-E). Therefore, Dga1 plays a major role in LD accumulation during chronic hyperosmotic stress, suggesting that LDs formed during chronic stress are regulated differently from those produced during the Lro1-dominated acute phase (Phan et al., 2025). This difference likely reflects distinct upstream cues and physiological functions for acute and chronic hyperosmotic stress.

Dga1 uses DAG and CoA-activated fatty acids to produce TAG (Figure S3A) (Oelkers et al., 2002). To further examine the role of Dga1 in LD accumulation during hyperosmotic stress, we restricted the availability of its substrates. Deletion of Nem1 or Faa1 and Faa4, which reduce the levels of DAG (Su et al., 2014) and activated fatty acids (Johnson et al., 1994), respectively, impaired LD accumulation under hyperosmotic stress (Figure S3B–J), consistent with a key role for Dga1 in this adaptive response.

Next, we examined Dga1 protein levels. Hyperosmotic stress resulted in a significant increase of endogenous Dga1 levels, as detected by western blot (Figure 2F,G). Importantly, this effect was specific as levels of all other neutral lipid–synthesizing enzymes remained unchanged in the same cells. The increase in Dga1 levels upon hyperosmotic stress was also detected by flow cytometry of cells expressing Dga1 endogenously tagged with mNeonGreen (Dga1-mNG) (Figure 2H). Using this approach, we also observed a modest increase in the levels of endogenous Lro1 expressed as a fusion to mCherry (Lro1-mCherry) while the levels of endogenous Are1 and Are2 remained unchanged (Figure S3K-M). The increase in Dga1-mNG levels under hyperosmotic stress was also confirmed by fluorescence microscopy. Interestingly, the increase in Dga1 levels and LD accumulation during hyperosmotic stress were accompanied by Dga1 redistribution from the ER to concentrate more prominently at LDs (Figure 2I). Together, these data highlight the central role of Dga1 in driving LD accumulation as part of the adaptive response to hyperosmotic stress.

### The canonical hyperosmotic stress response indirectly limits lipid droplet accumulation

Hyperosmotic stress adaptation is mediated through the HOG (High Osmolarity Glycerol) pathway (de Nadal and Posas, 2022). This well-characterized signalling cascade involves multiple sensors that converge on activation of the Hog1 MAP kinase (Figure S4A), which triggers an adaptive transcriptional program. Among other changes, this program rapidly enhances synthesis of glycerol, which acts as an intracellular osmolyte and facilitates restoration of cell volume (Figure 3A). Hog1 activation typically lasts for around 30 minutes, while downstream protein and glycerol synthesis peak within 60-90 minutes of stress onset (Petelenz-Kurdziel et al., 2013; Shen et al., 2023; Klipp et al., 2005). To test whether the LD accumulation upon hyperosmotic stress is under HOG pathway control, we analysed *hog1Δ* cells. In the absence of stress, *hog1Δ* cells showed Pln1-mCherry levels comparable to WT controls (Figure 3B). Remarkably, under hyperosmotic stress, *hog1Δ* cells displayed a dramatic increase in Pln1-mCherry levels, far exceeding that observed in WT cells (Figure 3B). This pronounced Pln1-mCherry increase in *hog1Δ* cells reflected the accumulation of more and larger LDs, as confirmed by fluorescence microscopy of BODIPY stained cells (Figure 3C-E). Moreover, manipulation of Hog1 activity by mutations that promote or inhibit its function resulted in a reduction or increase in Pln1-mCherry accumulation, respectively (Figure S4B-C). Together, these experiments indicate that the HOG pathway limits LD accumulation during hyperosmotic stress.

**Figure 3:**
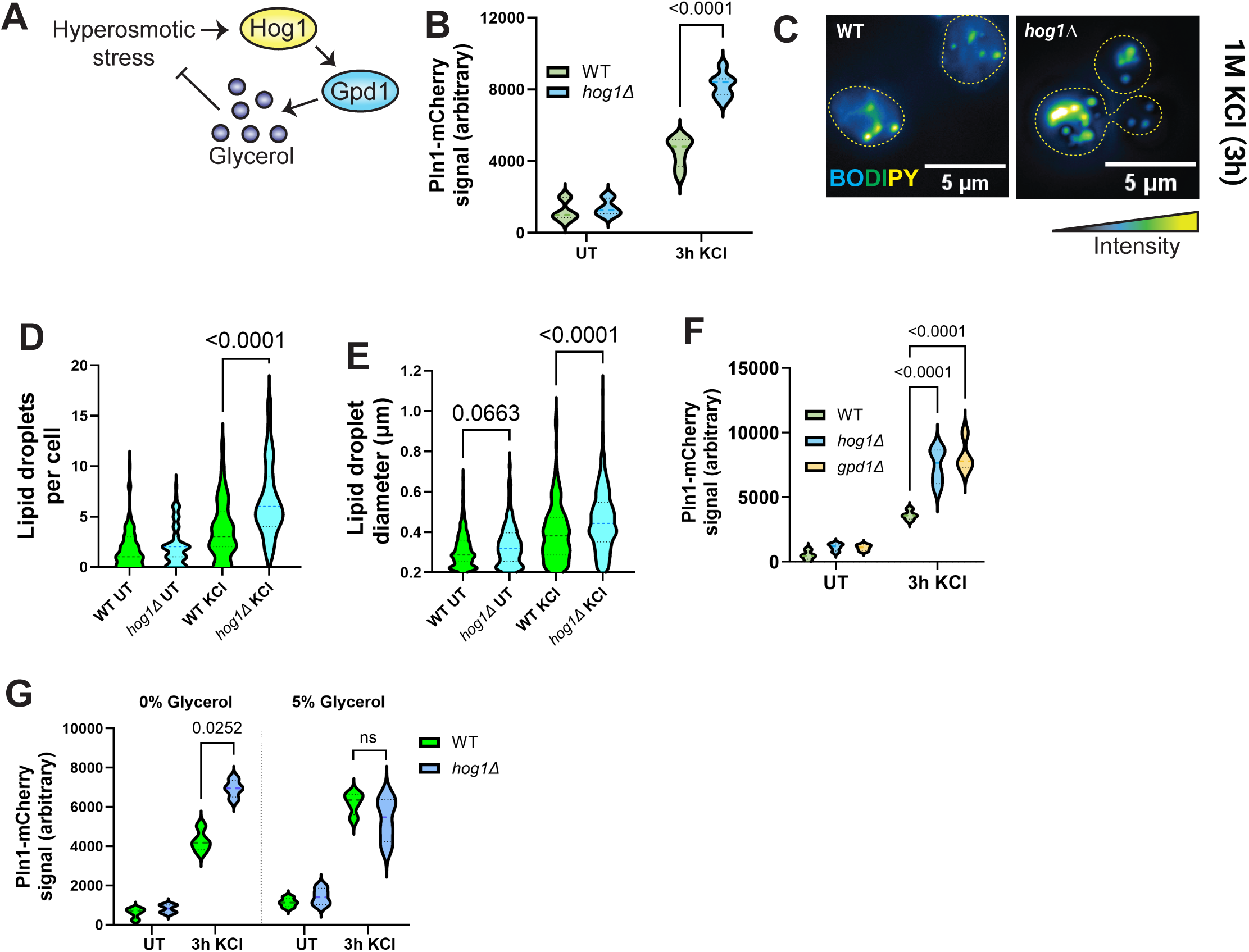
Canonical hyperosmotic stress signalling indirectly limits lipid droplet production. **A)** Simplified schematic of hyperosmotic stress adaptation signalling. Hyperosmotic stress leads to activation of the Hog1 MAP-kinase, which enhances Gpd1 production to create glycerol as a balancing intracellular osmolyte. **B)** Median Pln1-mCherry fluorescence measured by flow cytometry from populations of single wild-type (WT), and *hog1Δ* cells subjected to 1M KCl treatment in SCD media for 3 hours. UT = untreated. **C)** Lipid droplets stained with BODIPY and imaged by live cell microscopy after 3 hours 1M KCl treatment in SCD for Pln1-mCherry wild-type (WT) and *hog1Δ* cells. Z-projection images representative of multiple replicates. **D)** Analysis of lipid droplets per cell from samples used in (C). UT = untreated. **E)** Analysis of lipid droplet diameters from samples used in (C). UT = untreated. **F)** Median Pln1-mCherry fluorescence measured by flow cytometry from populations of single wild-type (WT), *hog1Δ*, and *gpd1Δ* cells subjected to 1M KCl treatment in SCD media for 3 hours. UT = untreated. **G)** Median Pln1-mCherry fluorescence measured by flow cytometry from populations of single wild-type (WT), and *hog1Δ* cells subjected to 1M KCl treatment with or without 5% glycerol in SCD media for 3 hours. UT = untreated.

The HOG pathway could limit LD accumulation because, when it is active, cells ultimately experience less of the hyperosmotic stress. In this scenario, one would expect that LD accumulation should be similarly modulated by altering glycerol levels. Indeed, deletion of *GPD1*, a glycerol phosphate dehydrogenase essential for glycerol production, enhanced Pln1-mCherry accumulation to the same extent as *HOG1* deletion (Figure 3F). Conversely, supplementing *hog1Δ* cells with glycerol reduced Pln1-mCherry accumulation to wild-type levels (Figure 3G). Blocking activation of the upstream sensor proteins Hkr1, Msb2 and Ypd1 did not prevent LD accumulation (Figure S4D), indicating that LD accumulation is not directly regulated by the HOG pathway. As cells defective in the HOG pathway cannot effectively produce the intracellular osmolytes necessary to resolve acute hyperosmotic stress (Petelenz-Kurdziel et al., 2013), LDs may be continually produced to counteract the stress. This could include helping to increase membrane tension, as described for the acute phase of hyperosmotic stress (Phan et al., 2025). Together, these results indicate that the HOG pathway indirectly limits LD accumulation by promoting cell adaptation to hyperosmotic stress.

### Chronic hyperosmotic stress drives membrane remodelling

Although our data identify TAG-containing LDs as important for adaptation to hyperosmotic stress, the broader consequences for cellular lipid composition remain unclear. We therefore performed whole-cell lipidomic analysis to define global lipid changes in response to chronic hyperosmotic stress. We observed that hyperosmotic stress resulted in a large increase of TAGs in WT and *hog1Δ* cells but not in *LDΔ*, as expected (Figure 4A). Interestingly, in *hog1Δ* cells hyperosmotic stress also resulted in a marked increase of STEs, while this was more modest in WT cells (Figure 4B). Thus, LDs produced in WT and *hog1Δ* cells upon chronic hyperosmotic stress appear to be compositionally distinct.

**Figure 4:**
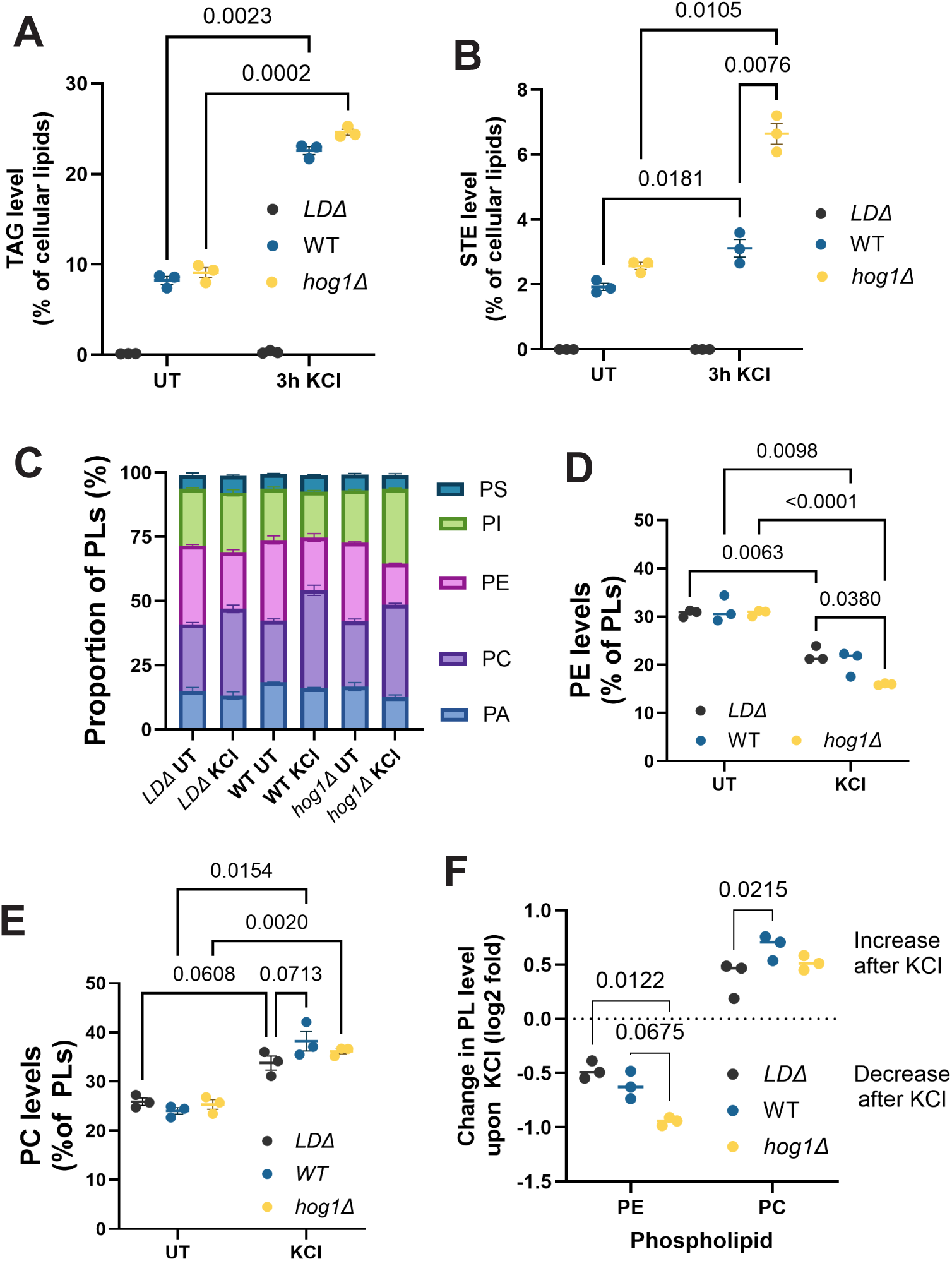
Chronic hyperosmotic stress drives lipid droplet-dependent lipidome changes. **A)** Triacylglycerols (TAGs) as a percentage of total cellular lipids detected with and without 3 hours of treatment with 1M KCl. **B)** Sterol esters (STEs) as a percentage of total cellular lipids detected with and without 3 hours of treatment with 1M KCl. **C)** Proportion of phospholipid species detected as a percentage of total phospholipids (PLs). PA = phosphatidic acid, PC = phosphatidylcholine, PE = phosphatidylethanolamine, PI = phosphatidylinositol, PS = phosphatidylserine. **D)** Phosphatidylethanolamine (PE) species as a percentage of total phospholipids (PLs) detected with and without 3 hours of treatment with 1M KCl. **E)** Phosphatidylcholine (PC) species as a percentage of total phospholipids (PLs) detected with and without 3 hours of treatment with 1M KCl. **F)** Log2 fold changes in PE and PC upon 1M KCl compared to untreated. Calculated from data in 4D and 4E.

We next examined changes to phospholipid composition. Most phospholipids remained largely unchanged as a proportion of the phospholipid pool (Figure 4C). Strikingly however, in all three strains analysed, hyperosmotic stress caused an increase in phosphatidylcholine (PC) levels with a concomitant drop in phosphatidylethanolamine (PE) levels (Figure 4D,E). The decrease in PE was comparable between WT and *LDΔ* cells, but more pronounced in *hog1Δ* cells, while the increase in PC was attenuated in the *LDΔ* cells compared to WT (Figure 4F). Thus, chronic hyperosmotic stress remodels the most abundant phospholipids and alters overall phospholipid composition.

### Interplay between lipid droplets and phospholipid remodelling during chronic hyperosmotic stress

To investigate the significance of PE/PC changes, we interfered with phospholipid synthesis. PE synthesis depends primarily on decarboxylation of PS by Psd1 and Psd2 in mitochondria and endosomes, respectively (Trotter et al., 1995; Gok et al., 2022) (Figure 5A). PC is produced by sequential tri-methylation of PE in the ER, with the initial step carried out by Cho2 and subsequent steps by Opi3 (McMaster and Bell, 1994) (Figure 5A). Both PE and PC can also be synthesised via the Kennedy pathway utilising DAG and either ethanolamine or choline, respectively. For PC synthesis via this pathway, the rate limiting step depends on Pct1 (Figure 5A).

**Figure 5:**
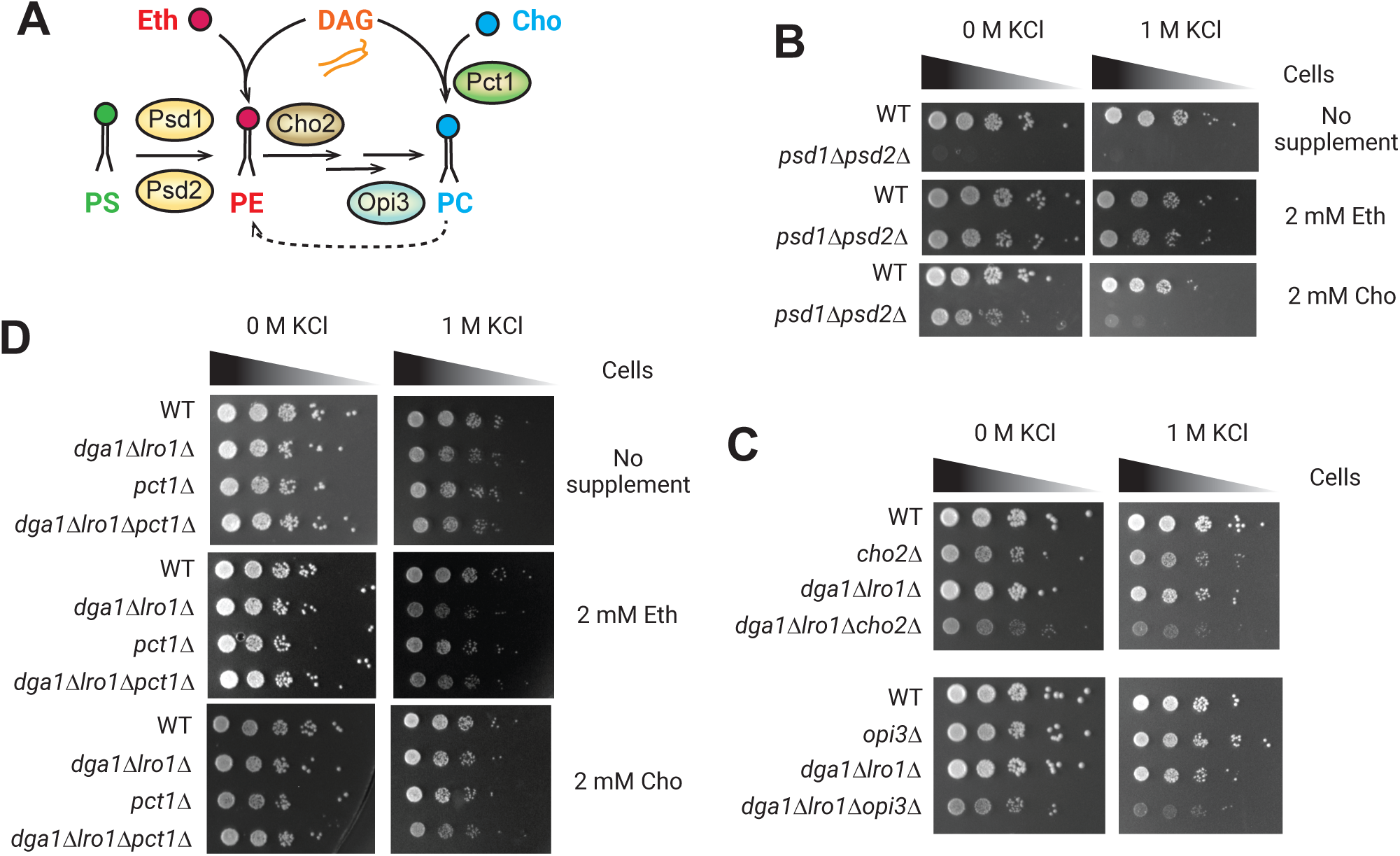
Phosphatidylethanolamine and phosphatidylcholine must be balanced for hyperosmotic stress adaptation. **A)** Schematic of phosphatidylethanolamine (PE) and phosphatidylcholine (PC) biosynthesis from phosphatidylserine (PS) conversion and combining diacylglycerol (DAG) with ethanolamine (Eth) or choline (Cho). **B)** Growth assay of wild-type (WT), *psd1Δpsd2Δ* and *cho2Δopi3Δ* cells after 2 days on SCD agar and SCD + 1M KCl agar with and without supplementation of 2mM ethanolamine (Eth) or choline (Cho). **C)** Growth assay of Pln1-mCherry wild-type (WT) and *dga1Δlro1Δ*, *cho2Δ*, *opi3Δ, dga1Δlro1Δcho2Δ*, and *dga1Δlro1Δopi3Δ* cells after 2 days on SCD agar and SCD + 1M KCl agar. **D)** Growth assay of Pln1-mCherry wild-type (WT) and *dga1Δlro1Δ*, *ptc1Δ,* and *dga1Δlro1Δpct1Δ* cells after 2 days on SCD agar and SCD + 1M KCl agar with and without supplementation of 2mM ethanolamine (Eth) or choline (Cho).

Given that both PE and PC are essential, the viability of *psd1Δpsd2Δ* cells depends on phospholipids synthesized via the Kennedy pathway and requires exogenous choline or ethanolamine (Trotter and Voelker, 1995; Iadarola et al., 2021; Storey et al., 2001). PE levels are reduced in these strains, even when supplemented with exogenous ethanolamine and choline (Trotter and Voelker, 1995). Choline supplementation of *psd1Δpsd2Δ* cells, only minimally suppressed sensitivity to hyperosmotic stress (Figure 5B), suggesting elevated PC alone is not sufficient for proper growth under this stress condition. Ethanolamine supplementation, which allows the synthesis of both PE and PC, largely rescued the growth defect (Figure 5B). Thus, while PE levels decrease upon hyperosmotic stress, a minimum PE level is nevertheless required for cell proliferation.

To assess the importance of increased PC levels under hyperosmotic stress, we interfered with its synthesis by deleting either *CHO2* or *OPI3* in *dga1Δlro1Δ* cells. In both cases, the mutations exacerbated the growth defect observed *dga1Δlro1Δ* cells (Figure 5D). The effect was specific for cells defective in TAG production, as neither *cho2Δ* nor *opi3Δ* mutants showed a growth defect upon hyperosmotic stress. Conversely, promoting PC synthesis by choline supplementation slightly improved the growth of *dga1Δlro1Δ* cells under hyperosmotic stress (Figure 5D and S5A,B). Importantly, this effect was dependent on Pct1, which is required for PC synthesis from exogenous choline (Figure 5D). Moreover, ethanolamine supplementation did not improve the growth of *dga1Δlro1Δ* cells under hyperosmotic stress. Thus, elevated PC and TAG levels are both important for adaptation to chronic hyperosmotic stress, and increasing PC levels can partially compensate for the absence of TAG.

### Upregulation of PC biosynthetic enzymes in the absence of TAG

Finally we investigated how hyperosmotic stress triggers an increase in PC levels. In particular, we tested whether this change is driven by upregulation of the key PC biosynthetic enzymes Cho2 and Opi3. Quantification of endogenously GFP-tagged Cho2 (Cho2-GFP) and Opi3 (Opi3-GFP) showed that chronic hyperosmotic stress increases the levels of both enzymes (Figure 6A,B). To further explore the links between TAG and PC synthesis in response to hyperosmotic stress, we tested how blocking TAG production impacts PC synthesis. Surprisingly, *dga1Δlro1Δ* cells displayed higher steady state levels of both Cho2-GFP and Opi3-GFP (Figure 6A,B). These levels did not increase further following hyperosmotic stress (Figure 6A,B). This effect appeared specific as other key enzymes in phospholipid metabolism were similarly responsive to hyperosmotic stress between the WT and mutant strains (Figure S5C,D). The increase in Cho2-GFP and Opi3-GFP levels in *dga1Δlro1Δ* cells could be due to changes in protein localization. However, both proteins retained a distribution consistent with ER localization, although some vacuolar fluorescence was observed for Opi3 (Figure 6C,D). To exclude potential confounding effects on flow cytometry measurements, we did Western blot analysis, which allows Opi3-GFP and free vacuolar GFP to be distinguished. This approach confirmed increased Opi3-GFP steady state levels both in WT cells under hyperosmotic stress and in TAG-deficient *dga1Δlro1Δ* cells (Figure S5E). Thus, loss of TAG synthesis leads to constitutive upregulation of PC biosynthetic enzymes.

**Figure 6:**
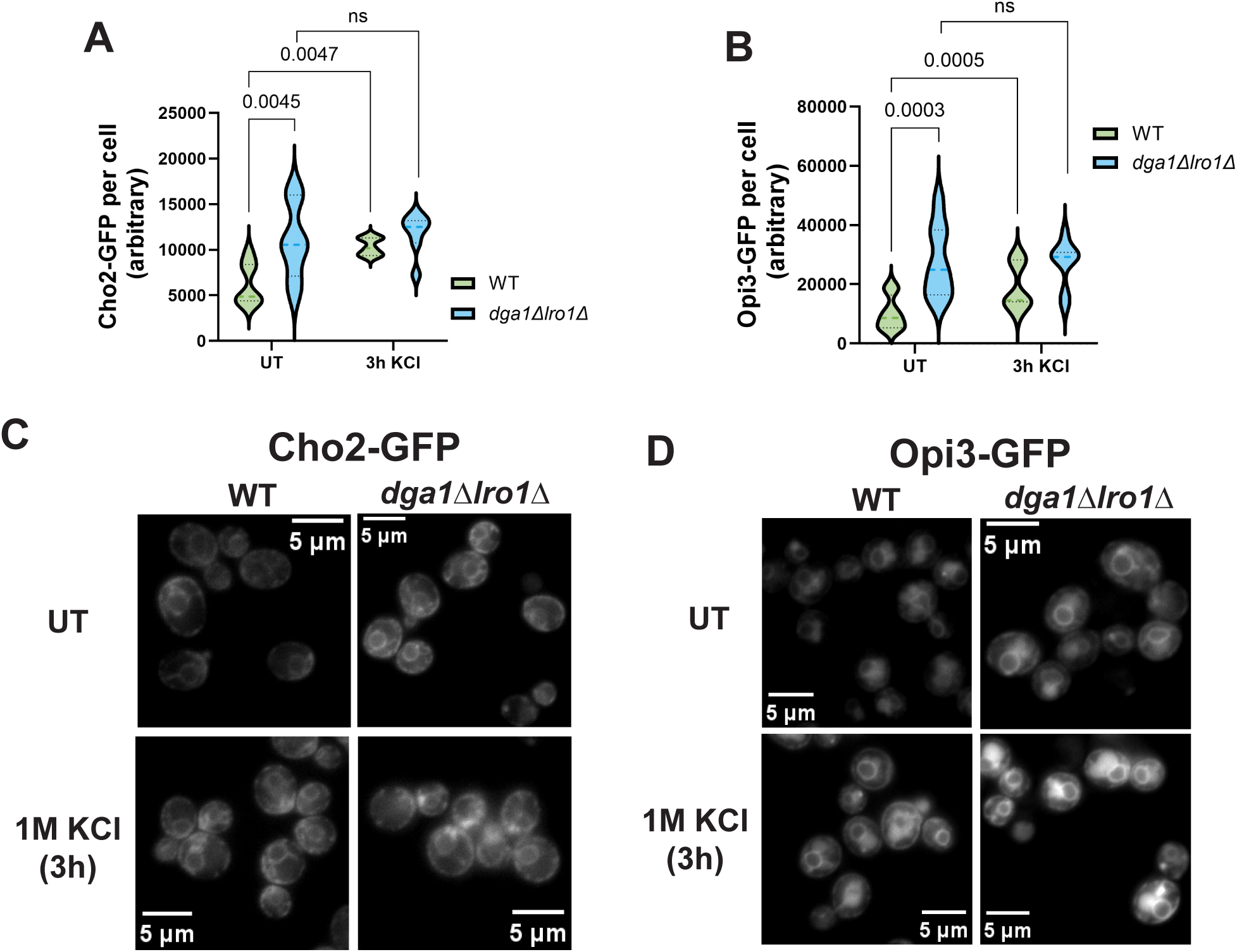
Phosphatidylcholine synthesis enzymes are misregulated in the absence of TAG. **A)** Median Cho2-GFP fluorescence measured by flow cytometry from populations of wild-type (WT) and *dga1Δlro1Δ* cells subjected to 1M KCl treatment in SCD media for 3 hours. UT = untreated. **B)** Median Opi3-GFP fluorescence measured by flow cytometry from populations of wild-type (WT) and *dga1Δlro1Δ* cells subjected to 1M KCl treatment in SCD media for 3 hours. UT = untreated. **C)** Localisation of Cho2 tagged with GFP in wild-type (WT) and *dga1Δlro1Δ* cells grown either in SCD media or SCD supplemented with 1M KCl for 3 hours prior to imaging. Images use the same colour-scale, show a single slice, and are representative of multiple experiments. **D)** Localisation of Opi3 tagged with GFP in wild-type (WT) and *dga1Δlro1Δ* cells grown either in SCD media or SCD supplemented with 1M KCl for 3 hours prior to imaging. Images use the same colour-scale, show a single slice, and are representative of multiple experiments.

## Discussion

We showed that chronic hyperosmotic stress triggers sustained and adaptive LD accumulation. Unlike the rapid and transient LD response observed during acute osmotic shock (Phan et al., 2025), chronic stress induces persistent LD accumulation that depends predominantly on Dga1-mediated TAG synthesis and is required for optimal growth. LD accumulation is not directly driven by HOG signalling, but instead becomes more prominent when osmotic balance is not fully restored. Thus, LD accumulation represents a parallel adaptive response to HOG signalling that supports optimal cell growth on hyperosmotic media.

Mechanistically, LD accumulation during chronic stress is distinct from the acute phase. Acute osmotic shock primarily engages Lro1-dependent TAG production (Phan et al., 2025), whereas chronic stress relies on Dga1. Although Dga1 and Lro1 are largely redundant for bulk TAG synthesis under steady-state conditions, only Dga1 is required for sustained LD accumulation during prolonged stress. This shift in enzymatic dependence suggests that chronic hyperosmotic stress activates a distinct lipid storage program, likely reflecting different physiological demands between transient membrane slack buffering and long-term membrane adaptation.

Our lipidomic profiling also revealed that chronic hyperosmotic stress is accompanied by extensive lipidome remodelling. We observe a shift in phospholipid composition, characterised by an elevation in PC and a concomitant reduction in PE. While the PE decrease occurs independently of LD formation, the magnitude of PC elevation is attenuated in cells lacking LDs. Genetic and metabolic manipulation of PC synthesis further demonstrates that increased PC contributes to fitness during chronic hyperosmotic stress, particularly when TAG synthesis is compromised. Together, these data support a model in which LD-driven TAG production and phospholipid remodelling are functionally coupled during adaptation.

The selective requirement for Dga1 may be particularly important in this context. Unlike Lro1, which can generate TAG by transferring acyl chains from membrane phospholipids (including PC), Dga1 utilises diacylglycerol and activated fatty acids (Oelkers et al., 2000). Preferential involvement of Dga1 during chronic stress may therefore promote TAG synthesis while preserving membrane phospholipid pools, facilitating coordinated lipid remodelling without depleting structural lipids. Consistent with this interpretation, TAG-deficient cells constitutively upregulate PC biosynthetic enzymes, yet remain compromised for growth under stress, suggesting that LD formation supports functional phospholipid adaptation beyond simple enzyme induction. How Dga1 and Opi3 are upregulated respectively during chronic hyperosmotic stress and TAG depletion remains unknown and should be addressed in future studies.

Why might increased PC levels be advantageous during chronic hyperosmotic stress? Hyperosmotic adaptation leads to sustained intracellular glycerol accumulation, which can directly interact with membrane headgroups and increase membrane rigidity (Abou-Saleh et al., 2019; Pocivavsek et al., 2011). Previous studies report reduced membrane fluidity under hyperosmotic conditions (Laroche et al., 2001). A shift from PE to PC may help counterbalance these effects. PE possesses a small, highly interactive headgroup that promotes tight packing and curvature stress, whereas PC has a bulkier, zwitterionic headgroup that favours bilayer stability and reduced curvature strain. Increasing PC levels may therefore buffer glycerol-induced rigidification and maintain membrane homeostasis during sustained hyperosmotic challenge.

An alternative, non-mutually exclusive possibility is that enhanced PC synthesis supports mitochondrial integrity. In yeast and mammalian systems, reduced PC levels impair mitochondrial architecture and function (Shiino et al., 2024; van der Veen et al., 2017; Schuler et al., 2016). Chronic hyperosmotic stress imposes metabolic and structural demands that may sensitize mitochondria to phospholipid imbalance (Criollo et al., 2007; Ikizawa et al., 2023). Increased PC production could therefore contribute to maintaining mitochondrial organisation during prolonged stress.

More broadly, accumulation of osmolytes such as glycerol is a conserved feature of hyperosmotic adaptation across eukaryotes (Burg and Ferraris, 2008). Salt tolerance in plants is enhanced by elevated PC levels, and is associated with TAG accumulation (Guo et al., 2019; Mueller et al., 2015; Corti et al., 2023). Renal epithelial cells, which experience chronic hyperosmotic stress, are enriched in both PC and TAG (Toback, 1984; Surma et al., 2021), and LD and PC accumulation are hallmarks of cancer cells subjected to mechanical and osmotic compression (Linke et al., 2024; Quan et al., 2025; Li et al., 2025). Our findings suggest that coordinated LD expansion and phospholipid remodelling may represent a conserved strategy for preserving membrane function during sustained osmotic imbalance.

## Materials and methods

### Yeast growth conditions

All strains used were derived from the BY4741 collection and are listed in Table S1. Mutants were made using either homologous recombination as described in detail elsewhere, or by crossing and dissection using a Singer Sporeplay+ microscope, followed by genotype determination. Strains which had not previously been verified were verified by performing PCRs within the deleted genes.

Cells were grown at 30°C on YPD agar plates (10 g/L yeast extract, 20 g/L peptone, 2% glucose, 20 g/L agar) before being used for the described experiments. For all growth assays, cells were grown in SCD (6.7 g/L YNB, 0.6 g/L CSM) in 16mm diameter glass tubes at 30°C, 180-200 rpm, for a minimum of 4 hours to allow them to enter logarithmic phase. For spot assays, they were then adjusted to 0.1 OD600 nm and serially diluted five times at a 1 in 10 ratio (such that each spot had one tenth the cells of the previous spot) before being spotted on the indicated plates and grown for 2 days at 30°C before imaging. For liquid growth assays, cells were diluted to 0.05 OD_600 nm_ in 200 ⍰l of the indicated media in a well of a flat-bottomed 96-well plate. The plate was incubated in a VantaStar (BMG Labtech) at 30°C for 18 hours, with 30 secs of vigorous shaking followed by OD_600 nm_ measurement every 15 minutes.

### Flow cytometry and analysis

Cells were grown to logarithmic phase the day before they were due to be analysed in SCD (16mm diameter glass tubes at 30°C, 180-200 rpm, for a minimum of 4 hours), unless otherwise indicated. Overnight they were diluted into 25mm diameter glass tubes or conical flasks, such that they would grow to ∼0.05-0.5 OD_600 nm_ after 16 hours at 30°C, 180-200 rpm. Samples were taken before treatment and stored on ice, then subjected to the indicated treatment with samples taken at the indicated times. These samples were then subject to flow cytometry analysis using a Fortessa (BD Biosciences) controlled by FACSDiva software (BD Biosciences) with a High-Throughput Sampler attachment set to record 10,000 events. Subsequent analysis was performed on FCS3.0 files using FlowJo (v10.10.0) to isolate bone fide single cells through gating, and the median relevant fluorescence value (either GFP or mCherry) for this population was recorded. From this median value, the median value of an unlabelled sample run alongside it was subtracted to obtain the final value.

### Microscopy and analysis

Cells were grown as for the flow cytometry experiments, with the exception that the overnight set-up aimed for the cells to grow to somewhere between 0.2-0.5 OD_600 nm_. Samples were taken at the indicated treatment times and labelled with either MDH (0.1 mM AutoDot, Abgent SM1000b) or BODIPY 493/503 (1 µg/ml, Invitrogen) and taken straight for imaging without washing. Imaging was performed on either a Zeiss AXIO Observer or Olympus FV3000 microscope with a 63x objective using Slidebook 6.0 (3i) or FV (Olympus) software respectively.

Analysis and image preparation was performed using FIJI. For analysis, images underwent Z-projection using the standard deviation setting. BODIPY stained lipid droplets were detected using ComDet (v.0.5.6) to isolate spherical particles and define their cross-sectional area in pixels, which was mathematically converted to diameter in Microsoft excel. Detected lipid droplets per cell were counted manually following ComDet particle identification, with budding cells counted as one cell if the bud was <50% of the size of the mother, and 2 separate cells if it was >50% the size of the mother. Any detected particle outwith a cell was excluded from further analysis.

### Protein extraction and Western blotting

Cells were grown as for the flow cytometry experiments, with the overnight growth occurring in 5-10 ml SCD in a conical flask. Cells were then diluted down to 0.2 OD_600 nm_ in SCD + 1M KCl, and samples of 2 OD600 nm harvested at the indicated times. UT samples were taken from the overnight culture prior to dilution. Samples were spun down in a benchtop centrifuge (8000rpm, 30sec) and resuspended in 300 µl ice-cold 150mM NaOH. After incubation for 10 minutes on ice, the cells were spun down again and resuspended in 100µl lysis buffer (0.1M Tris-HCl pH6.8, 2% SDS). Samples were incubated at 65°C with shaking in an Eppendorf Thermomixer F1.5 for 15 minutes, then brief spun to remove cell debris. Protein concentration of the supernatant was determined on a nanodrop, and the samples adjusted to the lowest value. 80µl of sample was then mixed with 20µl loading dye (50% glycerol, 0.05% bromophenol blue) and stored at −20°C prior to use.

For Western blot analysis, ∼100µg of protein samples were run on precast 4-20% TGX Criterion SDS-PAGE gels (Biorad) at 0.06 Amps per gel for 45 minutes. Blotting was performed using the TransBlot Turbo semi-dry system (Biorad) onto PVDF membranes. Gel running and blotting were performed according to manufacturer’s instructions. After blotting, membranes were blocked in 5% milk for 1 hour, then incubated overnight with primary antibodies. Following primary antibody incubation, secondaries were applied for at least one hour before imaging was performed using Western Lightening ECL Pro Enhanced Luminol Reagent (Revvity) on an Amersham Imager 600 (General Electric). Between each of these steps, membranes were twice rinsed with PBS-T (PBS with 0.1% Tween-20), followed by three 15-minute washes. Primary antibodies were all diluted in PBS-T 1% milk, 0.0002% sodium azide and stored at 4°C between uses. Secondary antibodies were diluted 1:10,000 in PBS-T and used once. Primary antibodies: α-Dga1 (1:3000, rabbit, Home-made), α-Flag (1:2000, mouse, Sigma-Aldrich A8592-2MG), α-HA (1:2000, rat, Roche, 3F10), α-Myc (1:1000, mouse, ThermoScientific 9E10), α-Tubulin (1:1000, rat, Santa Cruz, sc-53030), α-PGK1 (1:5000, mouse, ThermoFisher, 22C5D8).

Western blot analysis was performed using FIJI, with the average band intensity determined by drawing a box around it and using the measure feature. Intensity was normalised to that of the loading control (Tubulin or PGK1) to compensate for any errors in loading, then to the level of the untreated sample to account for differences between different membranes and imaging times.

### Lipidomic sample preparation

Cells were crashed out of frozen stocks onto a YPD agar plate and grown overnight at 30°C. Cells were then resuspended to ∼0.2 OD_600 nm_ in 2 ml SCD in a 16 mm diameter tube and grown for 4 hours at 30°C, 180 rpm. They were then diluted to 0.07 OD_600 nm_ in 5 ml SCD in a 25 mm diameter tube. After a further 4 hours growth at 30°C, 180 rpm they were diluted into 250ml SCD in a 1 litre conical flask so they would reach ∼0.4-0.8 OD_600 nm_ after 16 hours (wild-type and hog1Δ: to 0.0008 OD_600 nm_, are1Δare2Δdga1Δlro1Δ: to 0.001 OD600 nm). In the morning cells were inoculated in 250 ml SCD in 1 litre conical flasks at a concentration of 0.1 or 0.25 OD_600 nm_ for UT and KCl treated samples respectively and returned to the incubator. After 3 hours, equal OD_600 nm_ of all samples was taken and spun down using a JLA10.500 rotor (12 minutes, 4000rpm, 4°C). Pellets were resuspended in 10 ml ice-cold water, transferred to a 50 ml tube and spun down again using a benchtop centrifuge (5 minutes, 3900rpm, 4°C). The pellet was resuspended in 1 ml of ice-cold water, the OD_600 nm_ determined. 20 OD of cells transferred to individual pre-cooled 1.5 ml tubes. For all except 1 tube, these were spun down again and the pellets flash-frozen in liquid nitrogen before long-term storage at −70°C. For the other sample, the volume in the tube was made up to 500 µl by addition of ice-cold water and transferred to a 2 ml tube containing 0.5 ml 0.5 mm diameter glass beads (Thistle Scientific). Tubes were sealed with parafilm and subjected to lysis by bead beating at 4°C (OMNI International Bead Ruptor Elite, 10 x 2.9m/s 1 min on, 3 min off). After lysis, 0.5ml ice-cold water was added to the samples, mixed by pipetting, and 0.5ml was transferred without beads to a new 2 ml tube which was flash frozen in liquid nitrogen. Samples were stored at −70°C or on dry ice until they were ready to undergo analysis.

### Lipidomic analysis

Lipidomic mass spectrometry analysis was performed by Lipotype GmbH. Lipids were extracted using chloroform and methanol [Klose et al., 2012]. Samples were spiked with lipid class-specific internal standards prior to extraction. After drying and re-suspending in MS acquisition mixture, lipid extracts were subjected to mass spectrometric analysis. Mass spectra were acquired on a hybrid quadrupole/Orbitrap mass spectrometer equipped with an automated nano-flow electrospray ion source in both positive and negative ion mode. Lipid identification was performed using LipotypeXplorer [Herzog et al., 2011] on unprocessed (*.raw format) mass spectra. For MS-only mode, lipid identification was based on the molecular masses of the intact molecules. MSMS mode included the collision-induced fragmentation of lipid molecules and lipid identification was based on both the intact masses and the masses of the fragments. Prior to normalization and further statistical analysis lipid identifications were filtered according to mass accuracy, occupation threshold, noise and background. Lists of identified lipids and their intensities were stored in a database optimized for the particular structure inherent to lipidomic datasets. Intensity of lipid class-specific internal standards was used for lipid quantification.

### Statistical analysis

All statistical analysis and graph generation was performed using GraphPad Prism, with error bars representing the standard error. For dual-variable grouped graphs (Figures 2B, 3B, 3F, 3G, 4A, 4B, 4D, 4E, 4F, 6A, 6B, S1C-H, S2B, S2C, S2J, S3B, S3F, S3G, S4B-D, S5C-two way anova between matched replicates was performed, followed by multiple comparison analysis between samples of the same genotype and samples of the same treatment condition. For single-variable column graphs (Figures 1B, 1C, 1D, 1F, 2D, 2E, 3D, 3E, S1B, S2E, S2F, S2I, S2L, S2M, S3D, S3E, S3I, S3J), one-way anova with matched replicates was performed, followed by multiple comparisons between all samples. Paired t-tests were performed for Figures 2H, S3K-M. For Figure 2G a one sample t-test against an expected mean of 1 was performed.

## Data availability

All data used in this study are available upon request.

## Acknowledgments

We thank R. Hedley and V. Tsioligka at The Don Mason Facility of Flow Cytometry for technical assistance in the analysis of flow cytometry experiments, A. Wainman at the Light Microscopy Facility for help with imaging. We thank all members of the Carvalho lab for discussions and Z. Ji, L. Lemus and R. Klemm for critical reading of the manuscript. T.D. Williams was supported by BBSRC grant (BB/W001519/1) and by an Eric Reid Fund for Methodology grant awarded by the Biochemical Society. P. Carvalho was supported by a BBSRC grant (BB/W001519/1) and an investigator award from The Wellcome Trust (223153/Z/21/Z). The authors declare no competing financial interests.

## Author contributions

TDW performed most of the experiments; PC and TDW conceived the study, analyzed the data, and wrote the manuscript; all authors contributed to the final manuscript; PC supervised the project.

---

**Supplementary Table 1:** Strains used in this study.

| Reference | Genotype |
| --- | --- |
| yPC1505 | *MatA ura3Δ0 HIS33Δ1 leu2Δ0 met15Δ0* |
| yPC5042 | *MatA ura3Δ0 HIS33Δ1 leu2Δ0 met15Δ0 pln1-mCherry::HIS3* |
| yPC13064 | *Mat<sup>+</sup> ura3Δ0 HIS33Δ1 leu2Δ0 met15Δ0 pln1-mCherry::HIS3* |
| yPC5450 | *Mat<sup>+</sup> ura3Δ0 HIS33Δ1 leu2Δ0 met15Δ0 sei1Δ::NAT pln1-mCherry::HIS3* |
| yPC4929 | *Mat<sup>+</sup> ura3Δ0 HIS33Δ1 leu2Δ0 met15Δ0 are1Δ::KAN are2Δ::HYG dga1Δ::NAT lro1Δ::KAN* |
| yPC13088 | *Mat<sup>+</sup> ura3Δ0 HIS33Δ1 leu2Δ0 met15Δ0 lro1Δ::KAN pln1-mCherry::HIS3* |
| yPC13552 | *Mat<sup>+</sup> ura3Δ0 HIS33Δ1 leu2Δ0 met15Δ0 lro1Δ::KAN pln1-mCherry::HIS3* |
| yPC13538 | *Mat<sup>+</sup> ura3Δ0 HIS33Δ1 leu2Δ0 met15Δ0 cho2Δ::NAT psd1Δ::HYG psd2Δ::KAN pln1-mCherry::HIS3* |
| yPC13539 | *Mat<sup>+</sup> ura3Δ0 HIS33Δ1 leu2Δ0 met15Δ0 cho2Δ::NAT psd1Δ::HYG psd2Δ::KAN pln1-mCherry::HIS3* |
| yPC13082 | *Mat<sup>+</sup> ura3Δ0 HIS33Δ1 leu2Δ0 met15Δ0 are1Δ::KAN are2Δ::HYG pln1-mCherry::HIS3* |
| yPC13085 | *MatA ura3Δ0 HIS33Δ1 leu2Δ0 met15Δ0 dga1Δ::NAT lro1Δ::KAN pln1-mCherry::HIS3* |
| yPC13518 | *Mat<sup>+</sup> ura3Δ0 HIS33Δ1 leu2Δ0 met15Δ0 dga1Δ::NAT lro1Δ::KAN pln1-mCherry::HIS3* |
| yPC13565 | *Mat<sup>+</sup> ura3Δ0 HIS33Δ1 leu2Δ0 met15Δ0 dga1Δ::NAT pln1-mCherry::HIS3* |
| yPC13566 | *Mat<sup>+</sup> ura3Δ0 HIS33Δ1 leu2Δ0 met15Δ0 dga1Δ::NAT pln1-mCherry::HIS3* |
| yPC13356 | *Mat<sup>+</sup> ura3Δ0 HIS33Δ1 leu2Δ0 met15Δ0 are1-HA::HIS3 are2-MYC::KAN lro1-FLAG::HYG* |
| yPC10975 | *MatA ura3Δ0 HIS33Δ1 leu2Δ0 met15Δ0 dga1-mNeonGreen::HIS3* |
| yPC13068 | *Mat<sup>+</sup> ura3Δ0 HIS33Δ1 leu2Δ0 met15Δ0 nem1Δ::KAN pln1-mCherry::HIS3* |
| yPC13069 | *Mat<sup>+</sup> ura3Δ0 HIS33Δ1 leu2Δ0 met15Δ0 nem1Δ::KAN pln1-mCherry::HIS3* |
| yPC13342 | *MatA ura3Δ0 HIS33Δ1 leu2Δ0 met15Δ0 faa1Δ::KAN pln1-mCherry::HIS3* |
| yPC13343 | *MatA ura3Δ0 HIS33Δ1 leu2Δ0 met15Δ0 faa1Δ::KAN pln1-mCherry::HIS3* |
| yPC13496 | *Mat<sup>+</sup> ura3Δ0 HIS33Δ1 leu2Δ0 met15Δ0 faa4Δ::NAT pln1-mCherry::HIS3* |
| yPC13497 | *Mat<sup>+</sup> ura3Δ0 HIS33Δ1 leu2Δ0 met15Δ0 faa4Δ::NAT pln1-mCherry::HIS3* |
| yPC13376 | *Mat<sup>+</sup> ura3Δ0 HIS33Δ1 leu2Δ0 met15Δ0 faa1Δ::KAN faa4Δ::NAT pln1-mCherry::HIS3* |
| yPC13377 | *Mat<sup>+</sup> ura3Δ0 HIS33Δ1 leu2Δ0 met15Δ0 faa1Δ::KAN faa4Δ::NAT pln1-mCherry::HIS3* |
| yPC5609 | *Mat<sup>+</sup> ura3Δ0 HIS33Δ1 leu2Δ0 met15Δ0 lro1-mCherry::HIS3* |
| yPC6703 | *MatA ura3Δ0 HIS33Δ1 leu2Δ0 met15Δ0 are1-GFP::HIS5* |
| yPC5608 | *Mat<sup>+</sup> ura3Δ0 HIS33Δ1 leu2Δ0 met15Δ0 are2-mCherry::URA3* |
| yPC13120 | *Mat<sup>+</sup> ura3Δ0 HIS33Δ1 leu2Δ0 met15Δ0 hog1Δ::KAN pln1-mCherry::HIS3* |
| yPC13121 | *Mat<sup>+</sup> ura3Δ0 HIS33Δ1 leu2Δ0 met15Δ0 hog1Δ::KAN pln1-mCherry::HIS3* |
| yPC13206 | *Mat<sup>+</sup> ura3Δ0 HIS33Δ1 leu2Δ0 met15Δ0 hog1Δ::URA3 pln1-mCherry::HIS3* |
| yPC13196 | *Mat<sup>+</sup> ura3Δ0 HIS33Δ1 leu2Δ0 met15Δ0 gpd1Δ::KAN pln1-mCherry::HIS3* |
| yPC13197 | *Mat<sup>+</sup> ura3Δ0 HIS33Δ1 leu2Δ0 met15Δ0 gpd1Δ::KAN pln1-mCherry::HIS3* |
| yPC13245 | *Mat<sup>+</sup> ura3Δ0 HIS33Δ1 leu2Δ0 met15Δ0 hog1Δ::URA3 dga1Δ::NAT lro1Δ::KAN pln1-mCherry::HIS3* |
| yPC13249 | *Mat<sup>+</sup> ura3Δ0 HIS33Δ1 leu2Δ0 met15Δ0 hog1Δ::URA3 dga1Δ::NAT lro1Δ::KAN pln1-mCherry::HIS3* |
| yPC13244 | *Mat<sup>+</sup> ura3Δ0 HIS33Δ1 leu2Δ0 met15Δ0 hog1Δ::URA3 dga1Δ::NAT pln1-mCherry::HIS3* |
| yPC13247 | *Mat<sup>+</sup> ura3Δ0 HIS33Δ1 leu2Δ0 met15Δ0 hog1Δ::URA3 dga1Δ::NAT pln1-mCherry::HIS3* |
| yPC13246 | *Mat<sup>+</sup> ura3Δ0 HIS33Δ1 leu2Δ0 met15Δ0 hog1Δ::URA3 lro1Δ::KAN pln1-mCherry::HIS3* |
| yPC13248 | *Mat<sup>+</sup> ura3Δ0 HIS33Δ1 leu2Δ0 met15Δ0 hog1Δ::URA3 lro1Δ::KAN pln1-mCherry::HIS3* |
| yPC13203 | *Mat<sup>+</sup> ura3Δ0 HIS33Δ1 leu2Δ0 met15Δ0 sln1-DAmP::KAN pln1-mCherry::HIS3* |
| yPC13163 | *Mat<sup>+</sup> ura3Δ0 HIS33Δ1 leu2Δ0 met15Δ0 ptc1Δ::KAN pln1-mCherry::HIS3* |
| yPC13166 | *Mat<sup>+</sup> ura3Δ0 HIS33Δ1 leu2Δ0 met15Δ0 ptc1Δ::KAN pln1-mCherry::HIS3* |
| yPC13227 | *Mat<sup>+</sup> ura3Δ0 HIS33Δ1 leu2Δ0 met15Δ0 pbs2Δ::KAN pln1-mCherry::HIS3* |
| yPC13256 | *Mat<sup>+</sup> ura3Δ0 HIS33Δ1 leu2Δ0 met15Δ0 sho1Δ::HYG ssk1Δ::KAN pln1-mCherry::HIS3* |
| yPC13228 | *Mat<sup>+</sup> ura3Δ0 HIS33Δ1 leu2Δ0 met15Δ0 pbs2Δ::KAN pln1-mCherry::HIS3* |
| yPC13257 | *Mat<sup>+</sup> ura3Δ0 HIS33Δ1 leu2Δ0 met15Δ0 sho1Δ::HYG ssk1Δ::KAN pln1-mCherry::HIS3* |
| yPC13281 | *Mat<sup>+</sup> ura3Δ0 HIS33Δ1 leu2Δ0 met15Δ0 hkr1Δ::KAN msb2Δ::NAT pln1-mCherry::HIS3* |
| yPC13282 | *Mat<sup>+</sup> ura3Δ0 HIS33Δ1 leu2Δ0 met15Δ0 hkr1Δ::KAN msb2Δ::NAT pln1-mCherry::HIS3* |
| yPC13289 | *Mat<sup>+</sup> ura3Δ0 HIS33Δ1 leu2Δ0 met15Δ0 hkr1Δ::KAN ypd1-DAmP::NAT pln1-mCherry::HIS3* |
| yPC13290 | *Mat<sup>+</sup> ura3Δ0 HIS33Δ1 leu2Δ0 met15Δ0 hkr1Δ::KAN ypd1-DAmP::NAT pln1-mCherry::HIS3* |
| yPC13287 | *Mat<sup>+</sup> ura3Δ0 HIS33Δ1 leu2Δ0 met15Δ0 ypd1-DAmP::KAN msb2Δ::NAT pln1-mCherry::HIS3* |
| yPC13288 | *Mat<sup>+</sup> ura3Δ0 HIS33Δ1 leu2Δ0 met15Δ0 ypd1-DAmP::KAN msb2Δ::NAT* |
|  | *pln1-mCherry::HIS3* |
| yPC13585 | *MatA ura3Δ0 HIS33Δ1 leu2Δ0 met15Δ0 psd1-GFP::HIS3* |
| yPC13594 | *Mat? ura3Δ0 HIS33Δ1 leu2Δ0 met15Δ0 dga1Δ::NAT lro1Δ::KAN psd1-GFP::HIS3* |
| yPC13595 | *Mat? ura3Δ0 HIS33Δ1 leu2Δ0 met15Δ0 dga1Δ::NAT lro1Δ::KAN psd1-GFP::HIS3* |
| yPC13587 | *Mat? ura3Δ0 HIS33Δ1 leu2Δ0 met15Δ0 spo14-GFP::HIS3* |
| yPC13592 | *Mat? ura3Δ0 HIS33Δ1 leu2Δ0 met15Δ0 dga1Δ::NAT lro1Δ::KAN spo14-GFP::HIS3* |
| yPC13593 | *Mat? ura3Δ0 HIS33Δ1 leu2Δ0 met15Δ0 dga1Δ::NAT lro1Δ::KAN spo14-GFP::HIS3* |
| yPC13113 | *MatA ura3Δ0 HIS33Δ1 leu2Δ0 met15Δ0 hog1Δ::KAN* |
| yPC13516 | *Mat? ura3Δ0 HIS33Δ1 leu2Δ0 met15Δ0 nem1Δ::HYG dga1Δ::NAT lro1Δ::KAN pln1-mCherry::HIS3* |
| yPC13517 | *Mat? ura3Δ0 HIS33Δ1 leu2Δ0 met15Δ0 nem1Δ::HYG dga1Δ::NAT lro1Δ::KAN pln1-mCherry::HIS3* |
| yPC13285 | *Mat? ura3Δ0 HIS33Δ1 leu2Δ0 met15Δ0 ole1-DAmP::KAN pln1-mCherry::HIS3* |
| yPC13286 | *Mat? ura3Δ0 HIS33Δ1 leu2Δ0 met15Δ0 ole1-DAmP::KAN pln1-mCherry::HIS3* |
| yPC13212 | *Mat? ura3Δ0 HIS33Δ1 leu2Δ0 met15Δ0 spt23Δ::KAN pln1-mCherry::HIS3* |
| yPC13213 | *Mat? ura3Δ0 HIS33Δ1 leu2Δ0 met15Δ0 spt23Δ::KAN pln1-mCherry::HIS3* |
| yPC13210 | *Mat? ura3Δ0 HIS33Δ1 leu2Δ0 met15Δ0 mga2Δ::KAN pln1-mCherry::HIS3* |
| yPC13211 | *Mat? ura3Δ0 HIS33Δ1 leu2Δ0 met15Δ0 mga2Δ::KAN pln1-mCherry::HIS3* |
| yPC13529 | *Mat? ura3Δ0 HIS33Δ1 leu2Δ0 met15Δ0 mga2Δ::HYG dga1Δ::NAT lro1Δ::KAN pln1-mCherry::HIS3* |
| yPC13530 | *Mat? ura3Δ0 HIS33Δ1 leu2Δ0 met15Δ0 mga2Δ::HYG dga1Δ::NAT lro1Δ::KAN pln1-mCherry::HIS3* |
| yPC13507 | *Mat? ura3Δ0 HIS33Δ1 leu2Δ0 met15Δ0 psd1Δ::HYG psd2Δ::KAN pln1-mCherry::HIS3* |
| yPC13508 | *Mat? ura3Δ0 HIS33Δ1 leu2Δ0 met15Δ0 psd1Δ::HYG psd2Δ::KAN pln1-mCherry::HIS3* |
| yPC13460 | *Mat? ura3Δ0 HIS33Δ1 leu2Δ0 met15Δ0 opi3Δ::HYG cho2Δ::KAN pln1-mCherry::HIS3* |
| yPC13461 | *Mat? ura3Δ0 HIS33Δ1 leu2Δ0 met15Δ0 opi3Δ::HYG cho2Δ::KAN pln1-mCherry::HIS3* |
| yPC13544 | *Mat? ura3Δ0 HIS33Δ1 leu2Δ0 met15Δ0 pct1Δ::HYG pln1-mCherry::HIS3* |
| yPC13553 | *Mat? ura3Δ0 HIS33Δ1 leu2Δ0 met15Δ0 pct1Δ::HYG dga1Δ::NAT lro1Δ::KAN pln1-mCherry::HIS3* |
| yPC13554 | *Mat? ura3Δ0 HIS33Δ1 leu2Δ0 met15Δ0 pct1Δ::HYG dga1Δ::NAT lro1Δ::KAN pln1-mCherry::HIS3* |
| yPC13523 | *Mat? ura3Δ0 HIS33Δ1 leu2Δ0 met15Δ0 cho2Δ::HYG pln1-mCherry::HIS3* |
| yPC13534 | *Mat? ura3Δ0 HIS33Δ1 leu2Δ0 met15Δ0 cho2Δ::HYG dga1Δ::NAT lro1Δ::KAN pln1-mCherry::HIS3* |
| yPC13535 | *Mat? ura3Δ0 HIS33Δ1 leu2Δ0 met15Δ0 cho2Δ::HYG dga1Δ::NAT lro1Δ::KAN pln1-mCherry::HIS3* |
| yPC13441 | *Mat? ura3Δ0 HIS33Δ1 leu2Δ0 met15Δ0 opi3Δ::HYG pln1-mCherry::HIS3* |
| yPC13532 | *Mat? ura3Δ0 HIS33Δ1 leu2Δ0 met15Δ0 opi3Δ::HYG dga1Δ::NAT lro1Δ::KAN pln1-mCherry::HIS3* |
| yPC13533 | *Mat? ura3Δ0 HIS33Δ1 leu2Δ0 met15Δ0 opi3Δ::HYG dga1Δ::NAT lro1Δ::KAN pln1-mCherry::HIS3* |
| yPC6692 | *MatA ura3Δ0 HIS33Δ1 leu2Δ0 met15Δ0 cho2-GFP::HIS5* |
| yPC5926 | *MatA ura3Δ0 HIS33Δ1 leu2Δ0 met15Δ0 opi3-GFP::HIS5* |
| yPC13569 | *Mat? ura3Δ0 HIS33Δ1 leu2Δ0 met15Δ0 dga1Δ::NAT lro1Δ::KAN cho2-GFP::HIS5* |
| yPC13570 | *Mat? ura3Δ0 HIS33Δ1 leu2Δ0 met15Δ0 dga1Δ::NAT lro1Δ::KAN cho2-GFP::HIS5* |
| yPC13573 | *Mat? ura3Δ0 HIS33Δ1 leu2Δ0 met15Δ0 dga1Δ::NAT lro1Δ::KAN opi3-GFP::HIS5* |
| yPC13459 | *MatA ura3Δ0 HIS33Δ1 leu2Δ0 met15Δ0 spo14Δ::HYG pln1-mCherry::HIS3* |
| yPC13502 | *Mat? ura3Δ0 HIS33Δ1 leu2Δ0 met15Δ0 spo14Δ::HYG dga1Δ::NAT lro1Δ::KAN pln1-mCherry::HIS3* |
| yPC13503 | *Mat? ura3Δ0 HIS33Δ1 leu2Δ0 met15Δ0 spo14Δ::HYG dga1Δ::NAT lro1Δ::KAN pln1-mCherry::HIS3* |
| yPC13546 | *Mat? ura3Δ0 HIS33Δ1 leu2Δ0 met15Δ0 nte1Δ::HYG pln1-mCherry::HIS3* |
| yPC13557 | *Mat? ura3Δ0 HIS33Δ1 leu2Δ0 met15Δ0 nte1Δ::HYG dga1Δ::NAT lro1Δ::KAN pln1-mCherry::HIS3* |
| yPC13558 | *Mat? ura3Δ0 HIS33Δ1 leu2Δ0 met15Δ0 nte1Δ::HYG dga1Δ::NAT lro1Δ::KAN pln1-mCherry::HIS3* |
| yPC13545 | *Mat? ura3Δ0 HIS33Δ1 leu2Δ0 met15Δ0 plb1Δ::HYG pln1-mCherry::HIS3* |
| yPC13559 | *Mat? ura3Δ0 HIS33Δ1 leu2Δ0 met15Δ0 plb1Δ::HYG dga1Δ::NAT lro1Δ::KAN pln1-mCherry::HIS3* |
| yPC13560 | *Mat? ura3Δ0 HIS33Δ1 leu2Δ0 met15Δ0 plb1Δ::HYG dga1Δ::NAT lro1Δ::KAN pln1-mCherry::HIS3* |

## References

Athenstaedt, K. 2010. Neutral lipids in yeast: synthesis, storage and degradation. In Handbook of hydrocarbon and lipid microbiology. K.N. Timmis, editor. Springer Berlin Heidelberg, Berlin, Heidelberg. 471–480.

Cartwright, B.R., and J.M. Goodman. 2012. Seipin: from human disease to molecular mechanism. J. Lipid Res. 53:1042–1055. doi:10.1194/jlr.R023754.

Corti, E., S. Falsini, S. Schiff, C. Tani, C. Gonnelli, and A. Papini. 2023. Saline Stress Impairs Lipid Storage Mobilization during Germination in Eruca sativa. Plants. 12. doi:10.3390/plants12020366.

Criollo, A., L. Galluzzi, M.C. Maiuri, E. Tasdemir, S. Lavandero, and G. Kroemer. 2007. Mitochondrial control of cell death induced by hyperosmotic stress. Apoptosis. 12:3–18. doi:10.1007/s10495-006-0328-x.

Dubots, E., S. Cottier, M.-P. Péli-Gulli, M. Jaquenoud, S. Bontron, R. Schneiter, and C. De Virgilio. 2014. TORC1 regulates Pah1 phosphatidate phosphatase activity via the Nem1/Spo7 protein phosphatase complex. PLoS ONE. 9:e104194. doi:10.1371/journal.pone.0104194.

Ernst, R., C.S. Ejsing, and B. Antonny. 2016. Homeoviscous adaptation and the regulation of membrane lipids. J. Mol. Biol. 428:4776–4791. doi:10.1016/j.jmb.2016.08.013.

Farese, R.V., and T.C. Walther. 2025. Essential biology of lipid droplets. Annu. Rev. Biochem. 94:447–477. doi:10.1146/annurev-biochem-091724-013733.

Fei, W., G. Shui, Y. Zhang, N. Krahmer, C. Ferguson, T.S. Kapterian, R.C. Lin, I.W. Dawes, A.J. Brown, P. Li, X. Huang, R.G. Parton, M.R. Wenk, T.C. Walther, and H. Yang. 2011. A role for phosphatidic acid in the formation of “supersized” lipid droplets. PLoS Genet. 7:e1002201. doi:10.1371/journal.pgen.1002201.

Gao, Q., D.D. Binns, L.N. Kinch, N.V. Grishin, N. Ortiz, X. Chen, and J.M. Goodman. 2017. Pet10p is a yeast perilipin that stabilizes lipid droplets and promotes their assembly. J. Cell Biol. 216:3199–3217. doi:10.1083/jcb.201610013.

Giménez-Andrés, M., T. Emeršič, S. Antoine-Bally, J.M. D’Ambrosio, B. Antonny, J. Derganc, and A. Čopič. 2021. Exceptional stability of a perilipin on lipid droplets depends on its polar residues, suggesting multimeric assembly. eLife. 10. doi:10.7554/eLife.61401.

Gok, M.O., N.O. Speer, W.M. Henne, and J.R. Friedman. 2022. ER-localized phosphatidylethanolamine synthase plays a conserved role in lipid droplet formation. Mol. Biol. Cell. 33:ar11. doi:10.1091/mbc.E21-11-0558-T.

Grippa, A., L. Buxó, G. Mora, C. Funaya, F.-Z. Idrissi, F. Mancuso, R. Gomez, J. Muntanyà, E. Sabidó, and P. Carvalho. 2015. The seipin complex Fld1/Ldb16 stabilizes ER-lipid droplet contact sites. J. Cell Biol. 211:829–844. doi:10.1083/jcb.201502070.

Guo, Q., L. Liu, and B.J. Barkla. 2019. Membrane lipid remodeling in response to salinity. Int. J. Mol. Sci. 20. doi:10.3390/ijms20174264.

Hariri, H., S. Rogers, R. Ugrankar, Y.L. Liu, J.R. Feathers, and W.M. Henne. 2018. Lipid droplet biogenesis is spatially coordinated at ER-vacuole contacts under nutritional stress. EMBO Rep. 19:57–72. doi:10.15252/embr.201744815.

Hariri, H., N. Speer, J. Bowerman, S. Rogers, G. Fu, E. Reetz, S. Datta, J.R. Feathers, R. Ugrankar, D. Nicastro, and W.M. Henne. 2019. Mdm1 maintains endoplasmic reticulum homeostasis by spatially regulating lipid droplet biogenesis. J. Cell Biol. 218:1319–1334. doi:10.1083/jcb.201808119.

Heeren, J., and L. Scheja. 2021. Metabolic-associated fatty liver disease and lipoprotein metabolism. Mol. Metab. 50:101238. doi:10.1016/j.molmet.2021.101238.

Hosios, A.M., M.E. Wilkinson, M.C. McNamara, K.C. Kalafut, M.E. Torrence, J.M. Asara, and B.D. Manning. 2022. mTORC1 regulates a lysosome-dependent adaptive shift in intracellular lipid species. Nat. Metab. 4:1792–1811. doi:10.1038/s42255-022-00706-6.

Iadarola, D.M., A. Joshi, C.B. Caldwell, and V.M. Gohil. 2021. Choline restores respiration in Psd1-deficient yeast by replenishing mitochondrial phosphatidylethanolamine. J. Biol. Chem. 296:100539. doi:10.1016/j.jbc.2021.100539.

Ikizawa, T., K. Ikeda, M. Arita, S. Kitajima, T. Soga, H. Ichijo, and I. Naguro. 2023. Mitochondria directly sense osmotic stress to trigger rapid metabolic remodeling via regulation of pyruvate dehydrogenase phosphorylation. J. Biol. Chem. 299:102837. doi:10.1016/j.jbc.2022.102837.

Jin, Y., Y. Tan, J. Wu, and Z. Ren. 2023. Lipid droplets: a cellular organelle vital in cancer cells. Cell Death Discov. 9:254. doi:10.1038/s41420-023-01493-z.

Johnson, D.R., L.J. Knoll, D.E. Levin, and J.I. Gordon. 1994. Saccharomyces cerevisiae contains four fatty acid activation (FAA) genes: an assessment of their role in regulating protein N-myristoylation and cellular lipid metabolism. J. Cell Biol. 127:751–762. doi:10.1083/jcb.127.3.751.

Klemm, R.W., and P. Carvalho. 2024. Lipid droplets big and small: basic mechanisms that make them all. Annu. Rev. Cell Dev. Biol. 40:143–168. doi:10.1146/annurev-cellbio-012624-031419.

Klipp, E., B. Nordlander, R. Krüger, P. Gennemark, and S. Hohmann. 2005. Integrative model of the response of yeast to osmotic shock. Nat. Biotechnol. 23:975–982. doi:10.1038/nbt1114.

Klug, Y.A., J.V. Ferreira, and P. Carvalho. 2024. A unifying mechanism for seipin-mediated lipid droplet formation. FEBS Lett. 598:1116–1126. doi:10.1002/1873-3468.14825.

Lange, M., M. Wölk, V.W. Li, C.E. Doubravsky, J.M. Hendricks, S. Kato, Y. Otoki, B. Styler, S.L. Johnson, C.A. Harris, K. Nakagawa, I.F. Snodgrass, D. Kim, J.W. Newman, M. Fedorova, and J.A. Olzmann. 2025. FSP1-mediated lipid droplet quality control prevents neutral lipid peroxidation and ferroptosis. Nat. Cell Biol. 27:1902–1913. doi:10.1038/s41556-025-01790-y.

Lee, S.-J., J. Zhang, A.M.K. Choi, and H.P. Kim. 2013. Mitochondrial dysfunction induces formation of lipid droplets as a generalized response to stress. Oxid. Med. Cell. Longev. 2013:327167. doi:10.1155/2013/327167.

Linke, J.A., L.L. Munn, and R.K. Jain. 2024. Compressive stresses in cancer: characterization and implications for tumour progression and treatment. Nat. Rev. Cancer. 24:768–791. doi:10.1038/s41568-024-00745-z.

Liu, N., Y. Yun, Y. Yin, M. Hahn, Z. Ma, and Y. Chen. 2019. Lipid droplet biogenesis regulated by the FgNem1/Spo7-FgPah1 phosphatase cascade plays critical roles in fungal development and virulence in Fusarium graminearum. New Phytol. 223:412–429. doi:10.1111/nph.15748.

Li, L., Y. Huang, Y. Gui, W. Xiang, M. Yang, Y. Hou, and M. Peng. 2025. The role of phosphatidylcholine metabolism in tumors. Med. Oncol. 42:450. doi:10.1007/s12032-025-03017-4.

Madeira, J.B., C.A. Masuda, C.M. Maya-Monteiro, G.S. Matos, M. Montero-Lomelí, and B.L. Bozaquel-Morais. 2015. TORC1 inhibition induces lipid droplet replenishment in yeast. Mol. Cell. Biol. 35:737–746. doi:10.1128/MCB.01314-14.

Markgraf, D.F., R.W. Klemm, M. Junker, H.K. Hannibal-Bach, C.S. Ejsing, and T.A. Rapoport. 2014. An ER protein functionally couples neutral lipid metabolism on lipid droplets to membrane lipid synthesis in the ER. Cell Rep. 6:44–55. doi:10.1016/j.celrep.2013.11.046.

Mathiowetz, A.J., and J.A. Olzmann. 2024. Lipid droplets and cellular lipid flux. Nat. Cell Biol. 26:331–345. doi:10.1038/s41556-024-01364-4.

McMaster, C.R., and R.M. Bell. 1994. Phosphatidylcholine biosynthesis via the CDP-choline pathway in Saccharomyces cerevisiae. Multiple mechanisms of regulation. J. Biol. Chem. 269:14776–14783. doi:10.1016/S0021-9258(17)36692-9.

Mueller, S.P., D.M. Krause, M.J. Mueller, and A. Fekete. 2015. Accumulation of extra-chloroplastic triacylglycerols in Arabidopsis seedlings during heat acclimation. J. Exp. Bot. 66:4517–4526. doi:10.1093/jxb/erv226.

de Nadal, E., and F. Posas. 2022. The HOG pathway and the regulation of osmoadaptive responses in yeast. FEMS Yeast Res. 22. doi:10.1093/femsyr/foac013.

Nagaraj, B., A.W. James, A. Mathivanan, and V. Nachiappan. 2023. Impairment of RPN4, a transcription factor, induces ER stress and lipid abnormality in Saccharomyces cerevisiae. Mol. Cell. Biochem. 478:2127–2139. doi:10.1007/s11010-022-04623-w.

Nguyen, T.B., and J.A. Olzmann. 2017. Lipid droplets and lipotoxicity during autophagy. Autophagy. 13:2002–2003. doi:10.1080/15548627.2017.1359451.

Oelkers, P., D. Cromley, M. Padamsee, J.T. Billheimer, and S.L. Sturley. 2002. The DGA1 gene determines a second triglyceride synthetic pathway in yeast. J. Biol. Chem. 277:8877–8881. doi:10.1074/jbc.M111646200.

Oelkers, P., A. Tinkelenberg, N. Erdeniz, D. Cromley, J.T. Billheimer, and S.L. Sturley. 2000. A lecithin cholesterol acyltransferase-like gene mediates diacylglycerol esterification in yeast. J. Biol. Chem. 275:15609–15612. doi:10.1074/jbc.C000144200.

Petelenz-Kurdziel, E., C. Kuehn, B. Nordlander, D. Klein, K.-K. Hong, T. Jacobson, P. Dahl, J. Schaber, J. Nielsen, S. Hohmann, and E. Klipp. 2013. Quantitative analysis of glycerol accumulation, glycolysis and growth under hyper osmotic stress. PLoS Comput. Biol. 9:e1003084. doi:10.1371/journal.pcbi.1003084.

Phan, J., M. Silva, R. Kohlmeyer, R. Ruethemann, L. Gay, E. Jorgensen, and M. Babst. 2025. Recovery of plasma membrane tension after a hyperosmotic shock. Mol. Biol. Cell. 36:ar45. doi:10.1091/mbc.E24-10-0436.

Pressly, J.D., M.Z. Gurumani, J.T. Varona Santos, A. Fornoni, S. Merscher, and H. Al-Ali. 2022. Adaptive and maladaptive roles of lipid droplets in health and disease. Am J Physiol, Cell Physiol. 322:C468–C481. doi:10.1152/ajpcell.00239.2021.

Quan, J., C. Zhang, X. Chen, X. Cai, and X. Luo. 2025. Lipid droplet - organelle crosstalk and its implication in cancer. Prog. Biophys. Mol. Biol. 197:11–20. doi:10.1016/j.pbiomolbio.2025.05.002.

Rambold, A.S., S. Cohen, and J. Lippincott-Schwartz. 2015. Fatty acid trafficking in starved cells: regulation by lipid droplet lipolysis, autophagy, and mitochondrial fusion dynamics. Dev. Cell. 32:678–692. doi:10.1016/j.devcel.2015.01.029.

Rao, M.J., B. Folger, J. Reus, A. Toulmay, J. Li, R. Zhang, W. Prinz, J. Goodman, and F. Wang. 2025. Pln1 Mediates Lipid Droplet-Vacuole Tethering During Microlipophagy in Saccharomyces cerevisiae. bioRxiv.

Rogers, S., H. Hariri, N.E. Wood, N.O. Speer, and W.M. Henne. 2021. Glucose restriction drives spatial reorganization of mevalonate metabolism. eLife. 10. doi:10.7554/eLife.62591.

Safi, R., P. Menéndez, and A. Pol. 2024. Lipid droplets provide metabolic flexibility for cancer progression. FEBS Lett. 598:1301–1327. doi:10.1002/1873-3468.14820.

Sandager, L., M.H. Gustavsson, U. Ståhl, A. Dahlqvist, E. Wiberg, A. Banas, M. Lenman, H. Ronne, and S. Stymne. 2002. Storage lipid synthesis is non-essential in yeast. J. Biol. Chem. 277:6478–6482. doi:10.1074/jbc.M109109200.

Schuler, M.-H., F. Di Bartolomeo, C.U. Mårtensson, G. Daum, and T. Becker. 2016. Phosphatidylcholine affects inner membrane protein translocases of mitochondria. J. Biol. Chem. 291:18718–18729. doi:10.1074/jbc.M116.722694.

Seo, A.Y., P.-W. Lau, D. Feliciano, P. Sengupta, M.A.L. Gros, B. Cinquin, C.A. Larabell, and J. Lippincott-Schwartz. 2017. AMPK and vacuole-associated Atg14p orchestrate μ-lipophagy for energy production and long-term survival under glucose starvation. eLife. 6. doi:10.7554/eLife.21690.

Shen, W., Z. Gao, K. Chen, A. Zhao, Q. Ouyang, and C. Luo. 2023. The regulatory mechanism of the yeast osmoresponse under different glucose concentrations. iScience. 26:105809. doi:10.1016/j.isci.2022.105809.

Shiino, H., S. Tashiro, M. Hashimoto, Y. Sakata, T. Hosoya, T. Endo, H. Kojima, and Y. Tamura. 2024. Chemical inhibition of phosphatidylcholine biogenesis reveals its role in mitochondrial division. iScience. 27:109189. doi:10.1016/j.isci.2024.109189.

Storey, M.K., K.L. Clay, T. Kutateladze, R.C. Murphy, M. Overduin, and D.R. Voelker. 2001. Phosphatidylethanolamine has an essential role in Saccharomyces cerevisiae that is independent of its ability to form hexagonal phase structures. J. Biol. Chem. 276:48539–48548. doi:10.1074/jbc.M109043200.

Surma, M.A., M.J. Gerl, R. Herzog, J. Helppi, K. Simons, and C. Klose. 2021. Mouse lipidomics reveals inherent flexibility of a mammalian lipidome. Sci. Rep. 11:19364. doi:10.1038/s41598-021-98702-5.

Su, W.-M., G.-S. Han, and G.M. Carman. 2014. Yeast Nem1-Spo7 protein phosphatase activity on Pah1 phosphatidate phosphatase is specific for the Pho85-Pho80 protein kinase phosphorylation sites. J. Biol. Chem. 289:34699–34708. doi:10.1074/jbc.M114.614883.

Teixeira, V., T.S. Martins, W.A. Prinz, and V. Costa. 2021. Target of rapamycin complex 1 (TORC1), protein kinase A (PKA) and cytosolic ph regulate a transcriptional circuit for lipid droplet formation. Int. J. Mol. Sci. 22. doi:10.3390/ijms22169017.

Thiam, A.R., and E. Ikonen. 2021. Lipid Droplet Nucleation. Trends Cell Biol. 31:108–118. doi:10.1016/j.tcb.2020.11.006.

Toback, F.G. 1984. Phosphatidylcholine metabolism during renal growth and regeneration. Am. J. Physiol. 246:F249–59. doi:10.1152/ajprenal.1984.246.3.F249.

Trotter, P.J., J. Pedretti, R. Yates, and D.R. Voelker. 1995. Phosphatidylserine decarboxylase 2 of Saccharomyces cerevisiáe. Cloning and mapping of the gene, heterologous expression, and creation of the null allele. J. Biol. Chem. 270:6071–6080. doi:10.1074/jbc.270.11.6071.

Trotter, P.J., and D.R. Voelker. 1995. Identification of a non-mitochondrial phosphatidylserine decarboxylase activity (PSD2) in the yeast Saccharomyces cerevisiae. J. Biol. Chem. 270:6062–6070. doi:10.1074/jbc.270.11.6062.

van der Veen, J.N., J.P. Kennelly, S. Wan, J.E. Vance, D.E. Vance, and R.L. Jacobs. 2017. The critical role of phosphatidylcholine and phosphatidylethanolamine metabolism in health and disease. Biochim. Biophys. Acta Biomembr. 1859:1558–1572. doi:10.1016/j.bbamem.2017.04.006.

Wang, C.-W., R.-H. Chen, and Y.-K. Chen. 2024. The lipid droplet assembly complex consists of seipin and four accessory factors in budding yeast. J. Biol. Chem. 300:107534. doi:10.1016/j.jbc.2024.107534.

Wang, S., F.-Z. Idrissi, M. Hermansson, A. Grippa, C.S. Ejsing, and P. Carvalho. 2018. Seipin and the membrane-shaping protein Pex30 cooperate in organelle budding from the endoplasmic reticulum. Nat. Commun. 9:2939. doi:10.1038/s41467-018-05278-2.

Zadoorian, A., X. Du, and H. Yang. 2023. Lipid droplet biogenesis and functions in health and disease. Nat. Rev. Endocrinol. 19:443–459. doi:10.1038/s41574-023-00845-0.

