## Supplementary material for "Lipid remodelling enables adaptation to chronic hyperosmotic stress": Fig S1 to S4

Supplementary Figure 1

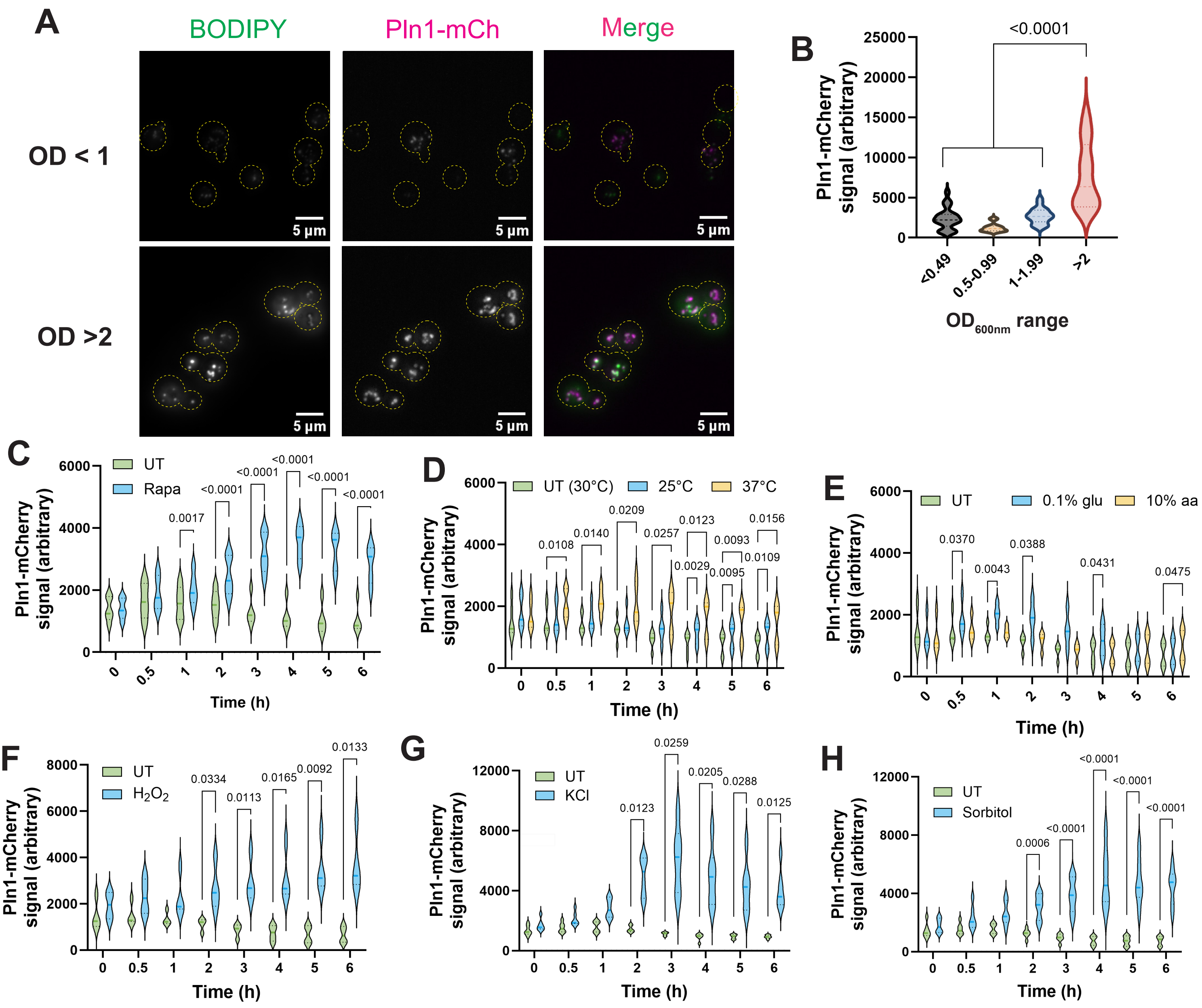

Supplementary Figure 2

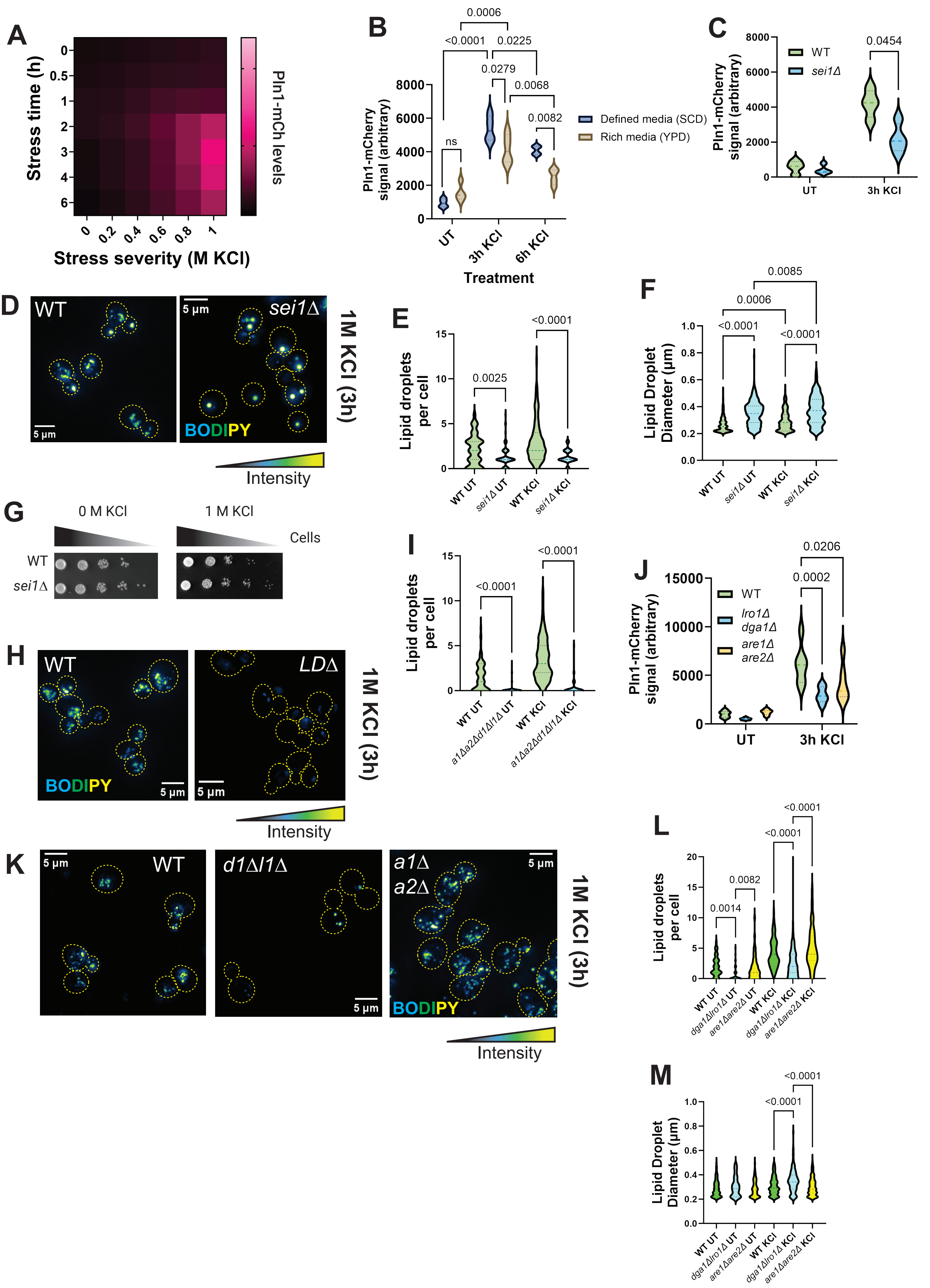

Supplementary Figure 3

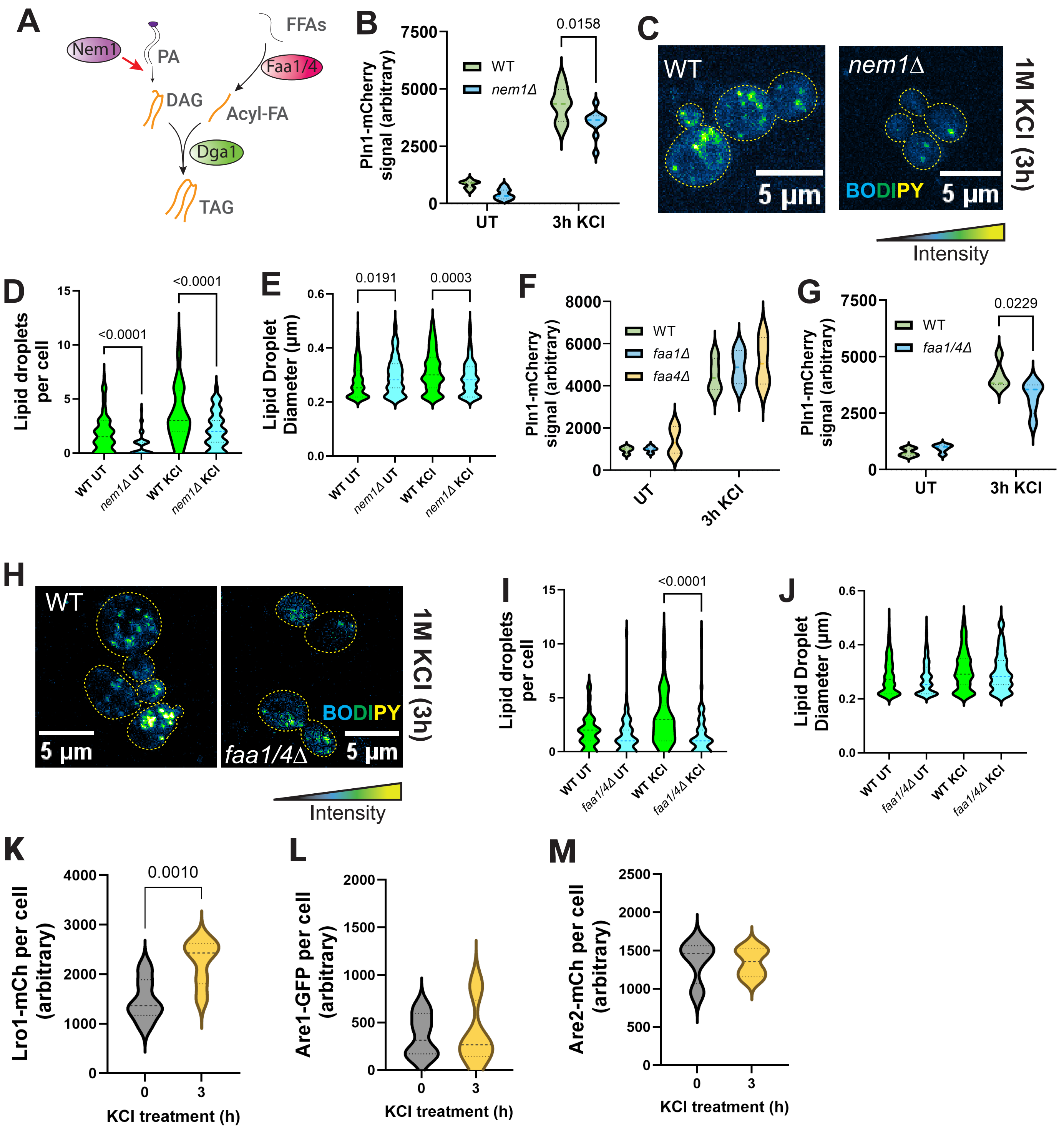

Supplementary Figure 4

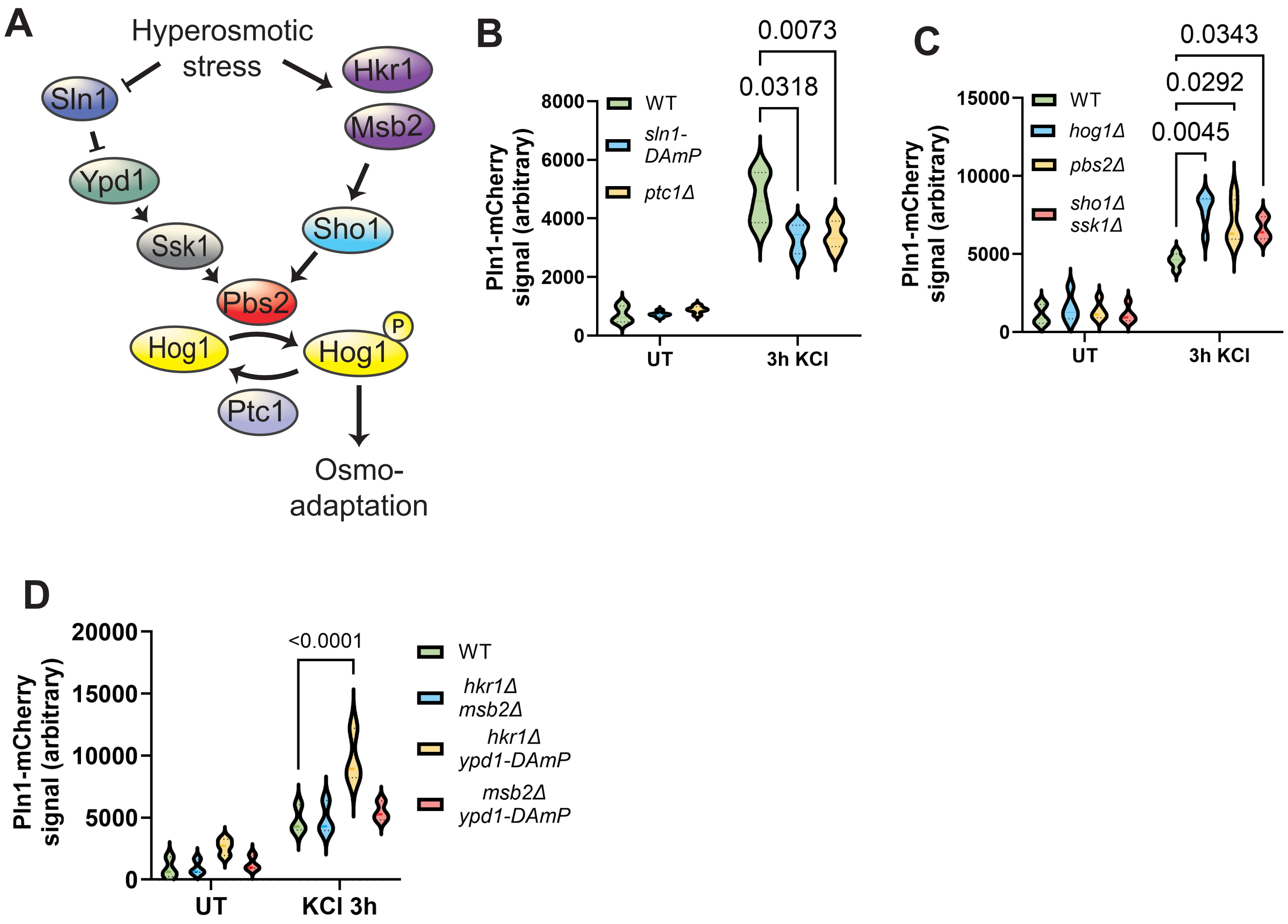

Supplementary Figure 5

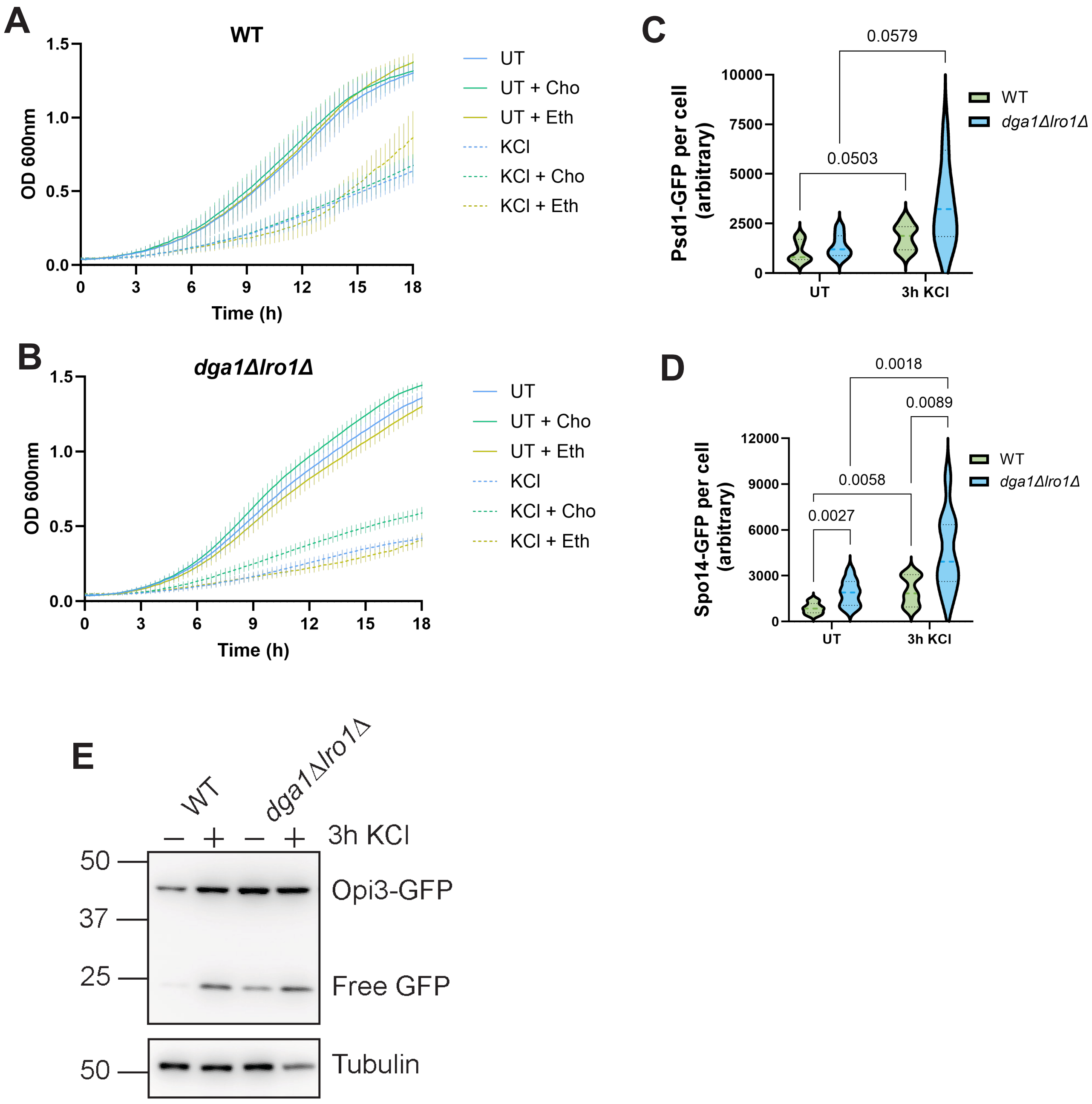
